# Ultrasensitive response in bacterial replication initiation

**DOI:** 10.64898/2026.05.28.728621

**Authors:** Alberto Sassi, Simone Pigolotti

**Affiliations:** Biological Complexity Unit, Okinawa Institute of Science and Technology, Graduate University, Onna, Okinawa 904-0495, Japan

## Abstract

In bacteria, the time at which genome replication initiates is carefully regulated, as precision in this timing is crucial for cell cycle stability. Theoretical and experimental studies have shown that the probability of replication initiation depends sharply on cell volume as the cell grows. Such sharp response is usually termed “ultrasensitive” in biology, and has been subject of extensive theoretical study. Nevertheless, the source of ultrasensitivity in this system remains unclear. In this work, we show how ultrasensitivity in replication initiation emerges from the dynamics of DnaA, the central protein regulating initiation in all bacterial species. To do so, we propose a mechanistic, thermodynamically consistent model of the dynamics of DnaA, and study the conditions under which an ultrasensitive response arises. We find that ultrasensitivity results from a combination of regulatory processes, that varies with growth conditions. In slow growth, we find an interplay between sequestration of DnaA on chromosomal binding sites and DnaA binding at the origin of replication. This mechanism requires a “sweet spot” of origin binding affinities, whose values are consistent with available experimental measurements. In fast growth, ultrasensitivity instead arises from a rapid transition between an inactive and an active form of DnaA, and requires cooperative binding at the origin of replication to be effective. Our results point to ultrasensitivity as the organizing principle underlying regulation of bacterial replication initiation.

## Introduction

Genome replication is a central event in the lifetime of a bacterial cell. The initiation of this process must be accurately timed to avoid instabilities in the cell cycle (***Wang and Levin, 2009; Scott et al., 2010; Mott and Berger, 2007; Tiruvadi-Krishnan et al., 2022; Zheng et al., 2016; Amir, 2014***). Initiation accuracy is especially important in bacteria such as *E. coli*, whose replication cycle lasts longer than the cell cycle in rich nutrient conditions (***Cooper and Helmstetter, 1968; Knöppel et al., 2023***). In this regime, multiple rounds of genome replication take place at the same time (***Cooper and Helmstetter, 1968; Fu et al., 2023; Ho and Amir, 2015; Knöppel et al., 2023; Zheng et al., 2016***), with multiple origins of replications firing within a narrow time window (***Boesen et al., 2024; Skarstad et al., 1986***). Empirical observations (***Wallden et al., 2016; Knöppel et al., 2023***) also show that the variability of cell volume at the time of initiation is small. Taken together, these observations imply that bacterial initiation is ultrasensitive, i.e., the origin firing probability must be an extremely steep function of the cell volume (***Berger and ten Wolde, 2023***).

Theoretical research has elucidated how an ultrasensitive response can arise in a biochemical system (***Ferrell and Ha, 2014a; Tu, 2008; Owen and Horowitz, 2023***). Possible mechanisms include binding cooperativity, allosteric change, titration and deactivation (***Ferrell and Ha, 2014a; Ferrell et al., 2014; Ferrell and Ha, 2014b; Buchler and Louis, 2008***). All of these mechanisms are indeed at play in bacterial replication initiation (***Kawakami et al., 2005; Kasho et al., 2017***). Specifically, initiation timing is mainly controlled by the protein DnaA (***Katayama et al., 1998; Flåtten et al., 2015; Knöppel et al., 2023; Li et al., 2025; Grant et al., 2011; Egan et al., 2004; Kaguni, 2006***). Before initiation, DnaA is progressively sequestered by strong binding sites (DnaA boxes) distributed along the chromosome; such sequestration acts as a titration mechanism (***Hansen et al., 1991; Olivi et al., 2025***). When these titration sites are close to saturation, DnaA binds cooperatively to weaker, clustered DnaA boxes located at the origin of replication (***Christensen et al., 1999; Speck et al., 1999; Schaper and Messer, 1995; Majka et al., 2001; Olivi et al., 2025***). This promotes allosteric melting of the double strand at the origin (***McGarry et al., 2004; Grimwade et al., 2018; Ozaki and Katayama, 2011***), thereby triggering initiation. Titration is a central aspect of the DnaA system, but it is not sufficient to ensure cell cycle stability in fast growth (***Berger and ten Wolde, 2022; Fu et al., 2023; Boesen et al., 2024; Knöppel et al., 2023***). Additionally, DnaA is present in the cell in ATP-bound and ADP-bound forms (***Katayama et al., 2017; Kasho et al., 2017; Knöppel et al., 2023; Fujimitsu et al., 2009***). Their interconversion provides an additional layer of regulation. Both forms can in principle melt the DNA double strand, but only DnaA-ATP is capable of cooperative binding at the origin (***Grimwade et al., 2018; McGarry et al., 2004; Kawakami et al., 2005***). Modeling shows that the dynamical regulation of the DnaA-ATP fraction is a stabilizing factor for the cell cycle in fast growth (***Berger and ten Wolde, 2022; Fu et al., 2023***).

In this paper, we develop a theory showing how these processes lead to an ultrasensitive response across growing conditions. We base our approach on a mechanistic, thermodynamically consistent model of DnaA kinetics coupled with the cell cycle. We show that this dynamics leads to an ultrasensitive response, quantified by very high values of the Hill coefficient, for physiologically relevant choices of parameters. In slow growth, sharp response arises when the origin of replication is bound at the time when progressive sequestration (titration) of DnaA on chromosomal binding sites is near completion. This condition sets precise constraints on the binding affinity of the origin sites, that are consistent with experimental measurements. In fast growth, ultrasensitivity stems from the interplay of fork-mediated activation/deactivation of DnaA near the time of initiation and origin binding. This mechanism exploits cooperative binding at the origin, as the ability to form cooperative interaction is the main distinction between DnaA-ATP and DnaA-ADP. Therefore, cooperative binding does not substantially contribute to ultrasensitivity per se as in classic biochemical models (***Hill, 1985***), but rather serves as a readout of the activation-deactivation switch. In fact, we show that cooperative binding is inessential in slow growth. Our theory rationalizes other aspects of the DnaA regulatory architecture in *E. coli*, such as the presence of strong binding sites at the origin of replication flanked by clusters of weak sites and sporadic clusters of binding sites far from the origin. Our results challenge existing paradigm for ultrasensitive response and show how the layered organization of the DnaA regulatory system guarantees precision in initiation across growth conditions.

## Results

### Equilibrium binding of DnaA along the cell cycle

Replication initiation is timed by titration in the following way. As the cell volume increases, the chromosomal binding sites are progressively diluted, while the concentration *a* of DnaA is kept constant by autorepression (***Braun et al., 1985; Saggioro et al., 2012; Iuliani et al., 2024; Ho and Amir, 2015***). DnaA thus first binds to chromosomal binding sites, and later to origin sites (Fig. 1a). Upon origin binding, initiation is triggered. Then, replication creates additional titration sites and DnaA is sequestered again, preventing further initiation events. After termination, the cycle starts again.

**Figure 1.**
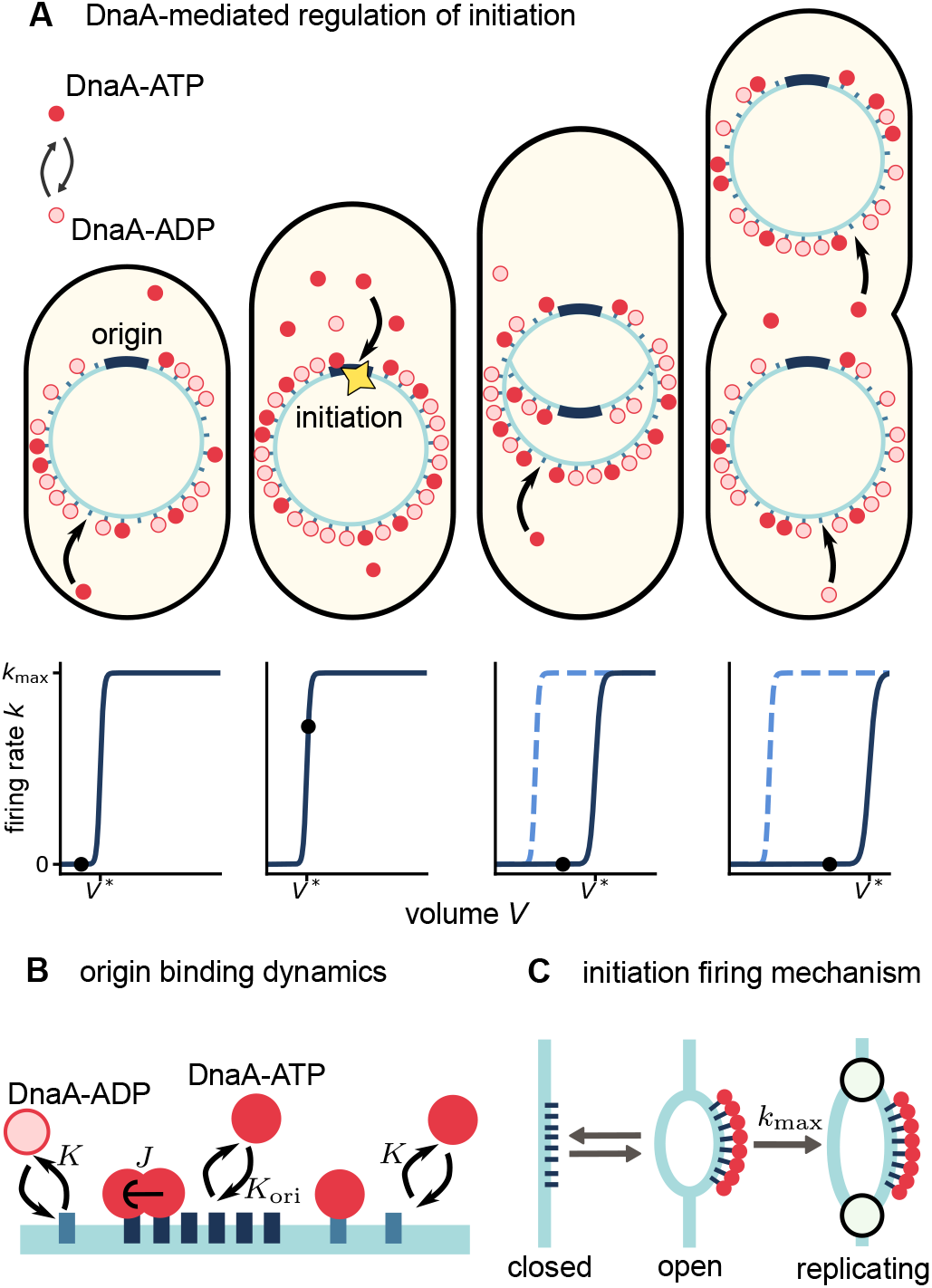
DnaA titration and replication initiation. (A) Titration mechanism. DnaA first binds to chromosomal sites. When they saturate, free DnaA starts to accumulate and bind to the weaker origin sites, thereby triggering initiation. After initiation, newly copied titration sites sequester DnaA again. During this cycle, the relative abundance of DnaA-ATP and DnaA-ADP is dynamically regulated. The outcome of this process is a steep (ultrasensitive) sigmoidal behavior of the firing rate as a function of the cell volume. The midpoint of the sigmoid shifts to larger volumes after the initiation time. (B) DnaA/DNA binding. DnaA binds monomerically to the titration sites (light blue) and cooperatively to the origin sites (dark blue). (C) Monod-Wyman-Changeux mechanism of replication initiation. DnaA binding stabilizes an open conformation at the origin of replication. The rate of initiation is proportional to the probability of open conformation.

The binding/unbinding of DnaA on DNA is extremely fast (∼ 10 − 200 ms) compared to the time scales of growth and protein synthesis (***Schenk et al., 2017; Olivi et al., 2025***). Consequently, we assume that DnaA/DNA binding is always at thermodynamic equilibrium (***Phillips, 2015***). Our model then includes two components: one describing equilibrium binding of DnaA and one describing the slower dynamical processes of cell growth, DNA replication, division, and regulation of the fraction of DnaA-ATP.

We assume that DnaA can bind to chromosomal and origin sites with binding constants *K* and *K*_ori_, respectively, with *K* ≪ *K*_ori_ (***Schaper and Messer, 1995; Majka et al., 2001; Fu et al., 2023***). These binding constants do not depend on the nucleotide state of DnaA. The origin sites are arranged in a cluster, to which DnaA-ATP proteins can bind cooperatively (***Keyamura et al., 2007; Kawakami et al., 2005; Noguchi et al., 2015; Majka et al., 2001***). This interaction is characterized by a free energy difference *J* (see Fig. 1b and Methods). The initiation rate *k* = *P*_open_*k*_max_ is proportional to the equilibrium probability *P*_open_ that the DNA at the origin is found in an open, partially melted conformation (Fig. 1c and (***Berger and ten Wolde, 2023***)). Here, the parameter *k*_max_ is the maximum initiation rate. By reducing free energy, binding of DnaA at the origin shifts the thermodynamic balance in favor of the open conformation – a mechanism analogous to the classic Monod-Wyman-Changeux model of allosteric change (***Monod et al., 1965***).

The dynamical component of the model includes growth, DNA replication, cell division, and interconversion between DnaA-ATP and DnaA-ADP (see Methods). We assume these dynamics to be deterministic for simplicity, aside for the initiation event that occurs stochastically.

### Multiple factors contribute to ultrasensitivity

Before initiation, the origin firing rate *k* increases with the volume *V* . Precise timing of initiation requires this rate to sharply transition from *k* ≈ 0 to *k* ≈ *k*_max_ at a certain volume *V* = *V* ^*^ (Fig. 2a). We describe this transition by a Hill function (***Mijatovi*ć *et al., 2026; Zhang et al., 2013***)

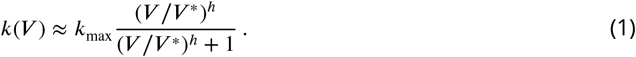

**Figure 2.**
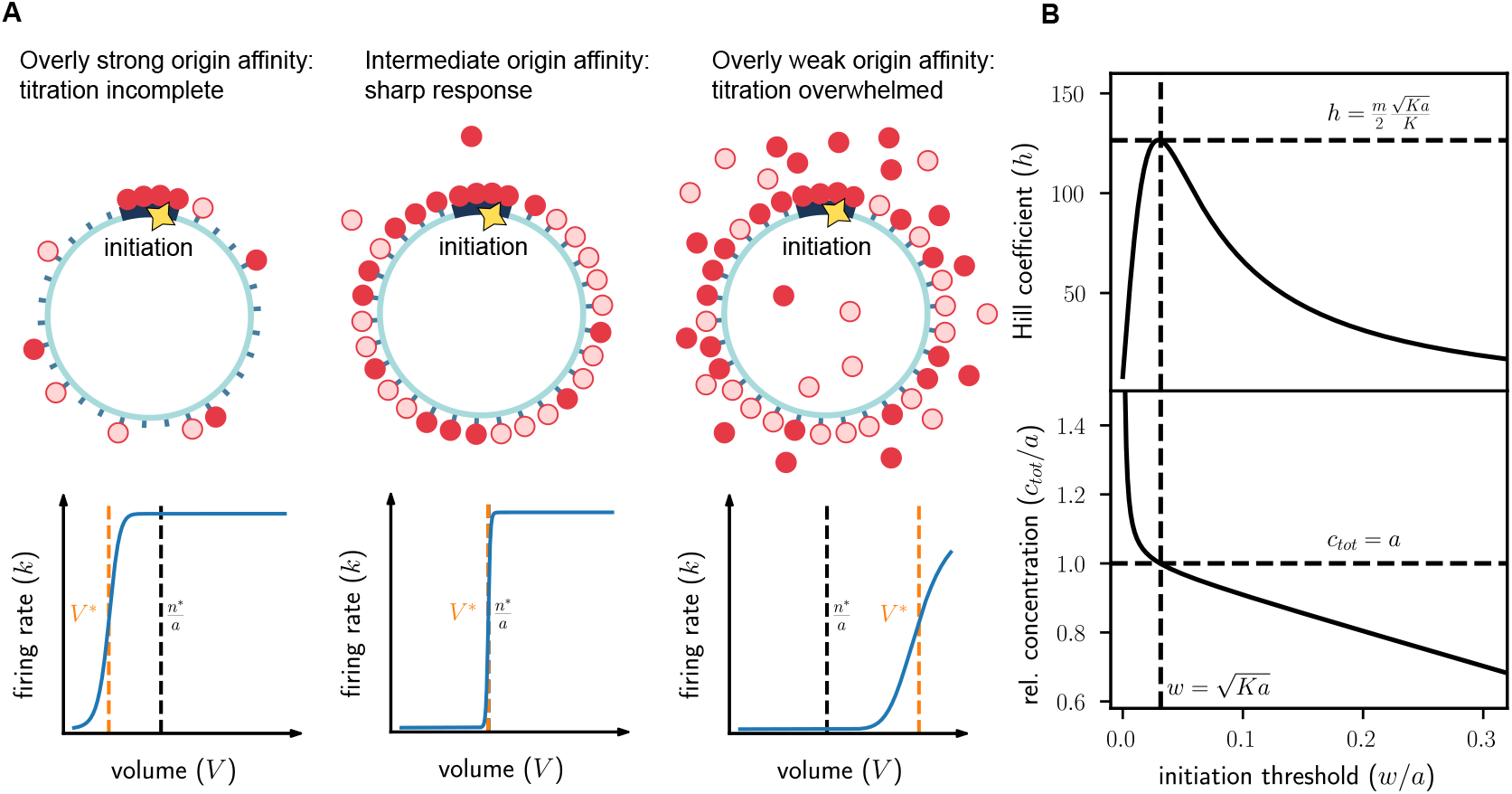
Ultrasensitivity requires tuning of binding parameters. (A) Effect of origin affinity on ultrasensitivity. To maximize the sharpening effect of titration, the effective affinity for the origin sites must be low enough that the titration sites are bound first, but it must be high enough that the origin sites are occupied as soon as the titration sites are close to saturation. Here *n*^*^ represents the total number of chromosomal sites at initiation. (B) Hill coefficient and relative concentration of titration sites. The Hill coefficient and the relative concentration of sites are plotted as a function of the relative concentration of free active DnaA at initiation *w*/*a*. The maximal value of the Hill coefficient is close to the point in which the concentrations of DnaA and titration sites are equal (dashed lines).

Ultrasensitivity amounts to having a large value of the Hill coefficient *h*.

To dissect the causes of sharp response, we decompose the Hill coefficient into three contribu-tions:

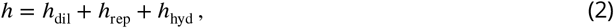

see Methods. The term *h*_dil_ represents the contribution to response due to dilution of chromosomal sites as the cell grows in volume. This term embodies the main effect of titration. The term *h*_rep_ is the contribution from creation of new titration sites by already present active forks in the multiround replication regime. This effect counteracts dilution by titration, so that *h*_dil_ > 0 while *h*_rep_ ≤ 0. The term *h*_hyd_ is the contribution to response from variations in the relative fraction of DNA-ATP. This effect enhances response sharpness (*h*_hyd_ ≥ 0), since the fraction of DnaA-ATP increases approaching initiation (see Methods), thereby facilitating precise timing (***Berger and ten Wolde, 2022***).

### Ultrasensitivity by titration requires tuning of origin affinity and cooperativity

We now focus on the slow growth regime, where *h*_rep_ = 0 before initiation, since active forks are absent. We assume that the main source of temporal regulation of the fraction of DnaA-ATP is fork-mediated hydrolysis. Therefore, the absence of forks also implies that *h*_hyd_ = 0 before initiation. We therefore have *h* = *h*_dil_ in this regime, i.e., the response is entirely due to dilution of titration sites.

We find that ultrasensitivity requires the origin sites to be occupied at the moment in which titration of chromosomal sites is complete (Fig. 2A). If origin sites are bound too strongly, initiation would take place before most titration sites are occupied. In this scenario, the buffering effect of chromosomal sites is not fully exploited. If origin sites are too weak, free DnaA accumulates in large amount before initiation is triggered. Also in this scenario, the effect of chromosomal sites is diminished, since a substantial amount of DnaA at the time of initiation is free rather than titrated.

We find that initiation takes place when the concentration of free DnaA-ATP reaches a threshold set by

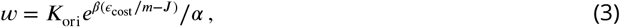

where *m* is the number of the origin binding sites, *ϵ*_cost_ is the intrinsic free energy difference associated with DNA melting at the origin, *α* = *a*_atp_/*a* with *a*_atp_ the concentration of DnaA-ATP, and β = 1/(*k*_B_*T* ) (see Methods and Appendix). We note that, since DnaA-ATP and DnaA-ADP are assumed to be equally titrated on the chromosome and origin sites are few compared with chromosomal sites, *α* also represents the fraction of free (cytoplasmic) DnaA-ATP over the total free DnaA.

The Hill coefficient is expressed by

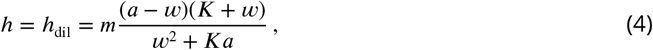

see Methods and Appendix. The function ( ) presents a maximum 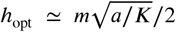 for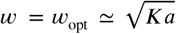 (Fig. 2B), where the approximation is valid for *K* ≪ *a*. This maximum embodies the optimal scenario of sharp response depicted in Fig. 2A. In fact, we find that at this maximum the concentration *a* of DnaA is equal to the total concentration *c*_tot_ of titration sites at the moment of initiation (Fig. 2B and Appendix). Since *K* ≪ *a*, this means that in the optimal case the titration sites are nearly saturated at the moment of initiation with little DnaA remaining free, consistently with the intuition anticipated in Fig. 2A.

Although cooperative binding is often associated with sharp response, we remark that in this case cooperativity at the origin is not necessary. In particular, the combination of parameters *w* in Eq. (3) can be tuned at its optimal value by setting *J* = 0 and appropriately choosing *K*_ori_.

### Replication and deactivation affect precision in opposing ways

In fast growth, all three contributions to the Hill coefficient in Eq. (2) should be taken into account. Mathematical expressions for *h*_rep_ and *h*_hyd_ (see Methods) depend on the model parameters, some of which are difficult to estimate experimentally. It is nevertheless useful to evaluate them for our estimated model parameters based on *E. coli* in slow and fast growth (see Table 1 in Methods). In slow growth, we obtain *h*_rep_ = *h*_hyd_ = 0 and we estimate *h* = *h*_dil_ ≈ 120, assuming optimal binding parameters (see Fig. 2B). In fast growth, we estimate that *h*_dil_ ≈ 100, *h*_rep_ ≈ −66 and *h*_hyd_ ≈ 45, so that *h* ≈ 79 (see Appendix). This estimate shows how, in fast growth, the increase in sharpness due to hydrolysis compensates for the loss due to creation of new titration sites before initiation.

**Table 1.** Parameters used in the titration-based replication model.

| Parameter | Description | Value |
| --- | --- | --- |
| $n_{\text{sites}}$ | Number of chromosomal titration sites per chromosome | 300 |
| $C$ | Replication time (min) | 40 |
| $D$ | Time between replication termination and division (min) | 20 |
| $\tau$ | Doubling time (min) | 25 (fast growth)<br>70 (slow growth) |
| $\Lambda$ | Growth rate | $\ln 2/\tau$ |
| $m$ | Number of weak origin sites | 8 |
| $\beta\epsilon_{\text{cost}}$ | Energetic cost of binding | 13 |
| $t_{\text{eclipse}}$ | Eclipse period (min) | 10 |
| $t_{\text{delay}}$ | Delay before initiation (min) | 2.5 |
| $a$ | Total DnaA concentration (nM) | 1000 |
| $K$ | Dissociation constant for strong sites (nM) | 1 |
| $\chi_0$ | Hydrolysis strength | 0.23 |
| $\alpha_{\text{max}}$ | Maximal active fraction | 1 (fast growth)<br>0.23 (slow growth) |
| $k_{\text{max}}$ | Maximum firing rate ($\text{min}^{-1}$) | 50 |

### Ultrasensitivity determines cell cycle stability

We now link sharpness of the response with the accuracy of the cell cycle, quantified by the coefficient of variation (CV) of the initiation volume. Empirically, this coefficient of variation is on the order of 10% for *E. coli* (***Wallden et al., 2016; Amir, 2014***). Achieving a small coefficient of variation requires a high value of the Hill coefficient (see Appendix). In slow growth, we find that accurate cell cycle timing requires the Hill coefficient to be close to its optimum value *h*_opt_ (Fig 3A). This supports that binding constants in *E*.*coli* must be close to optimality.

**Figure 3.**
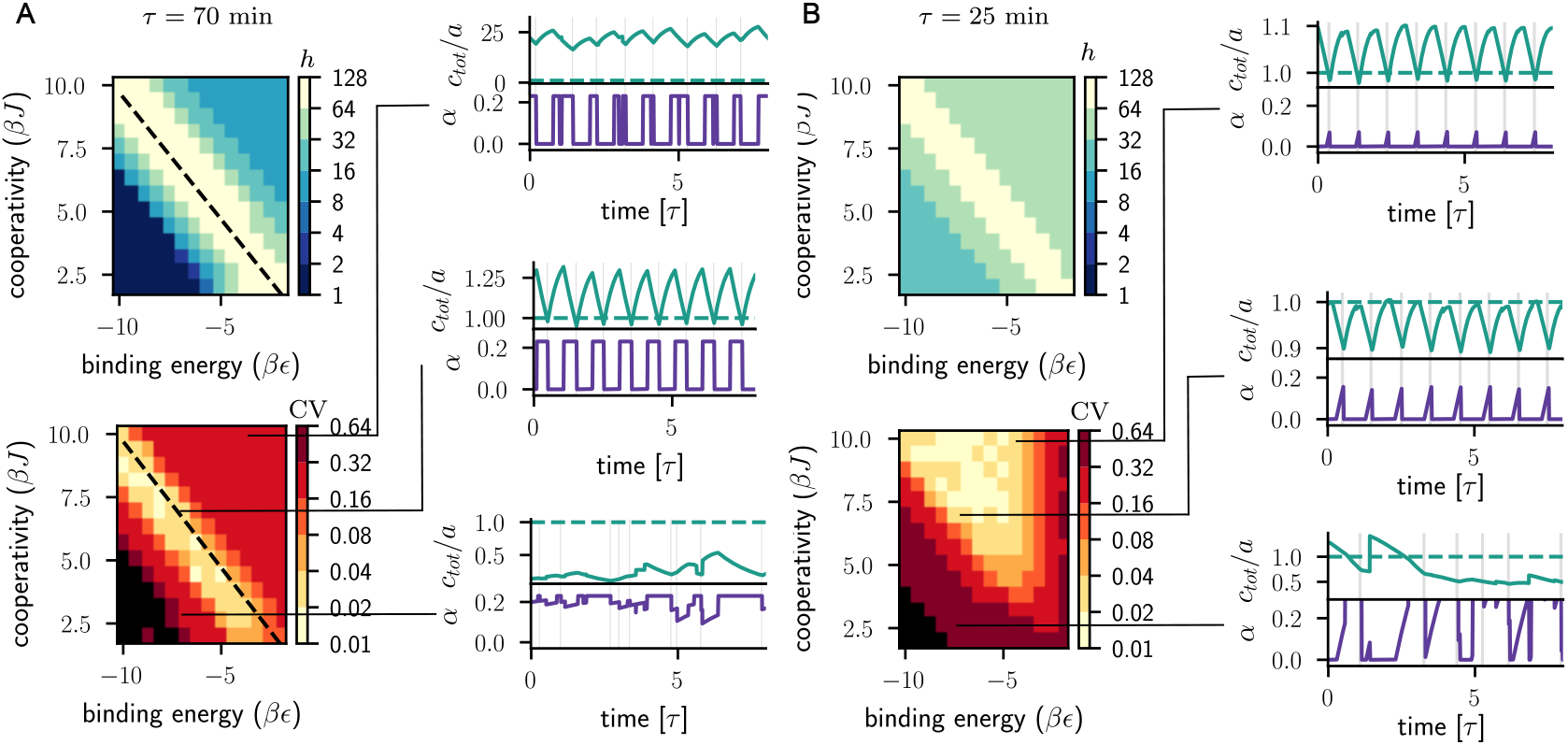
Different strategies ensure ultrasensitivity in slow and fast growth. Hill coefficient *h* and coefficient of variation (CV) of the initiation volume in slow *τ* = 70 min (A) and fast *τ* = 25 min (B) growth, as a function of binding energy ( *βϵ* = − ln[*vK*_ori]_, where = *v* 63nm^3^ is the excluded volume of a DnaA protein) and cooperativity. The black dashed line in (A) indicates the theoretical optimal condition 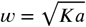. The black regions in the CV heatmaps denote a regime in which origin affinity is too low to trigger initiation for realistic volumes. Time traces (green) show the concentration of titration sites *c*_tot_ relative to the concentration of DnaA *a*. In the optimal case, this ratio is equal to one at the moment of initiation (green dashed lines). Active fraction of DnaA *α* are also shown (purple). In fast growth, the pulsatile dynamics of *α* is a main determinant of sharp response.

In fast growth, the activation/deactivation switch produces a pulsatile response in the active fraction *α* (Fig. 3B) which is more effective in maintaining timing than the square-wave response in slow growth (time traces in Fig. 3A). This pulsatile dynamics is caused by dilution of pre-existing active forks. In this respect, fork-mediated hydrolisis of Dna-ATP can be seen as a nonlinear sensor of the concentration of active forks. This nonlinearity is pronounced due to our assumption that fork-mediated hydrolysis operates close to saturation (see Appendix). This dynamics leads to a value of *h*_hyd_ sufficiently large to guarantee an ultrasensitive response outside of the optimal regime of binding parameters for the titration mechanism (upper right regions of the heatmaps in Fig. 3B). In this region, both *h*_dil_ and *h*_rep_ are small in absolute value. This means that the system is less reliant on the origin binding parameters and more on the activation/deactivation switch, in line with previous work (***Berger and ten Wolde, 2022***).

Since ultrasensitivity in fast growth depends on interconversion between DnaA-ATP and DnaA-ADP, cooperativity is crucial in this regime to discriminate between the two forms of DnaA. Consequently, the system is unstable if *J* is too low (Fig. 3B and (***Fu et al., 2023***)). We present a stability analysis characterizing this region in Appendix.

### Asymmetric chromosomal titration of active DnaA limits early re-initation events

Besides response sharpness, the DnaA system must be also effective at preventing reinitiation events. Reinitiation is partially inhibited by the fact that, shortly after initiation, the hemimethylated origin is not accessible to DnaA (***Lu et al., 1994; von Freiesleben et al., 2000***). This mechanism acts on a timescale *t*_eclipse_ of about 10 min; this time should be long enough so that all parallel origins are guaranteed to fire in fast growth (***Berger and ten Wolde, 2023***). To prevent later reinitiation, the initiation volume *V* ^*^ must be shifted at larger values after initiation (Fig. 1A). This shift is achieved by creation of new titration sites and fork-mediated deactivation of DnaA-ATP.

In this section, we discuss an additional mechanism that could contribute to an effective deactivation. So far, we have assumed for simplicity that binding at chromosomal sites is indifferent to the nucleotide state of DnaA. However, recent spt-PALM experiments show that the chromosomal occupancy is higher in mutant strains with impaired deactivation of DnaA-ATP (***Olivi et al., 2025***). A possible explanation is that, even far from the origin, cooperative binding allows DnaA-ATP to access clusters of sites that would not be accessible to DnaA-ADP, in line with recent findings (***Stringer et al., 2024***). To rationalize the role of these sites, we introduce an extension of our model in which the chromosome includes ordinary sites, that can be bound by either DnaA-ATP or DnaA-ADP, and a relative fraction *θ* of special sites, that can only be bound by DnaA-ATP (see Fig.4A). Both ordinary and special sites are assumed to be homogeneously distributed along the DNA, and are characterized by the same dissociation constant *K*.

We find that the special sites contribute to avoid early reinitiation events by selectively titrating DnaA-ATP after initiation. An analysis of the sigmoidal firing rate curve before and after the moment of initiation (Fig. 4B) shows that the rate is reduced more drastically in the presence of special sites. In particular, for small θ, the shift Δ*V* ^*^ in the midpoint volume increases in both *θ* and Δ*α*, where Δ*α* is the variation in the active fraction before and after initiation (Fig. 4C).

**Figure 4.**
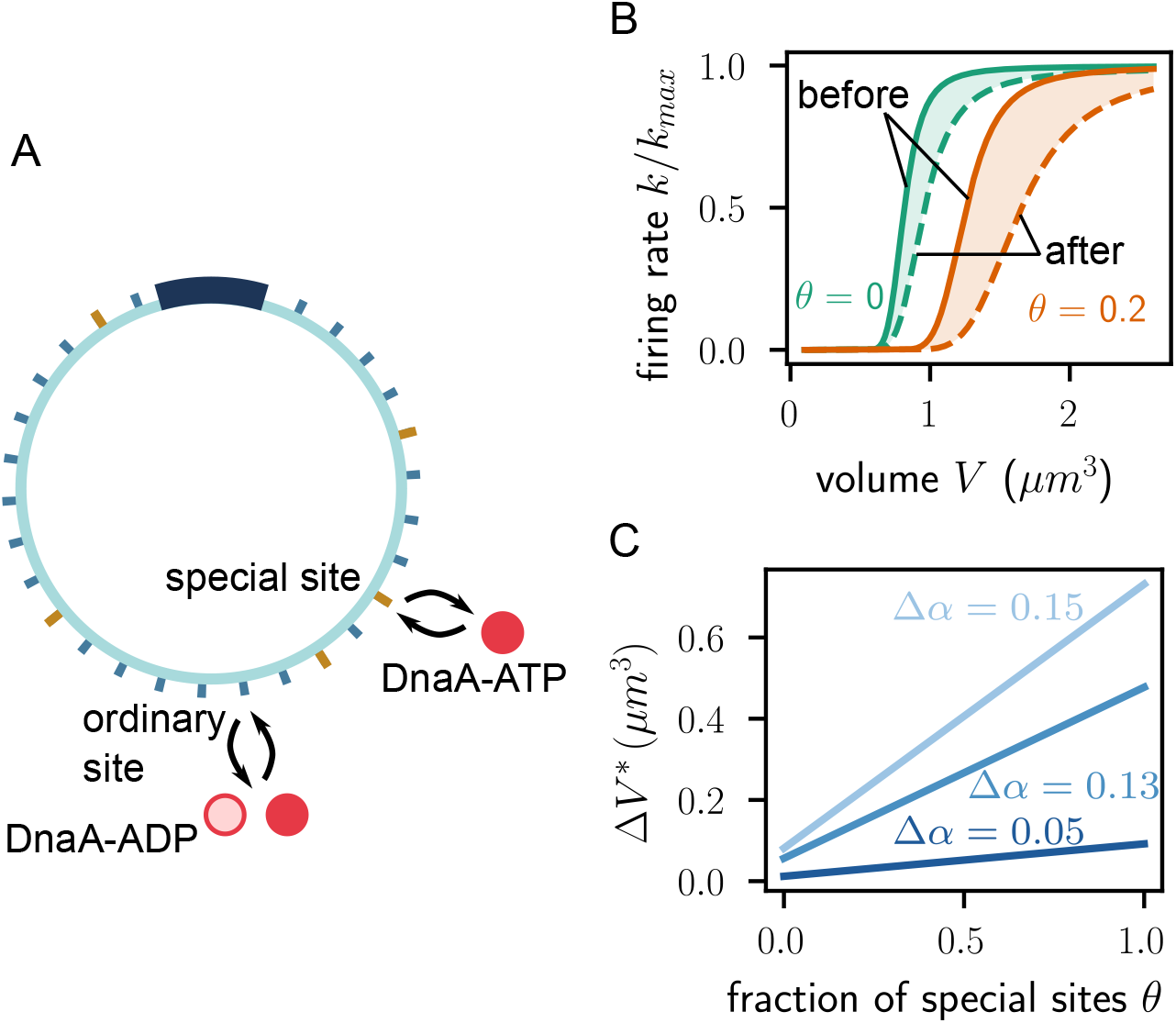
Asymmetric chromosomal binding of active and inactive DnaA. (A) A variant of the model in which a fraction *θ* of “special” chromosomal sites are only accessible to DnaA-ATP, while the rest of the sites can be bound by DnaA in both nucleotide states. (B) Firing rate as a function of the cell volume before (continuous line) and after (dashed line) replication initiation. Special sites shift the value of the initiation volume *V* ^*^ and make post-initiation deactivation more effective. (C) Shift in the initiation volume as a function of *θ* and the change in active fraction upon initiation Δ*α*. We note that, when *θ* > 0, the fraction of free DnaA-ATP differs from the fraction of total DnaA-ATP.

**Figure 5.**
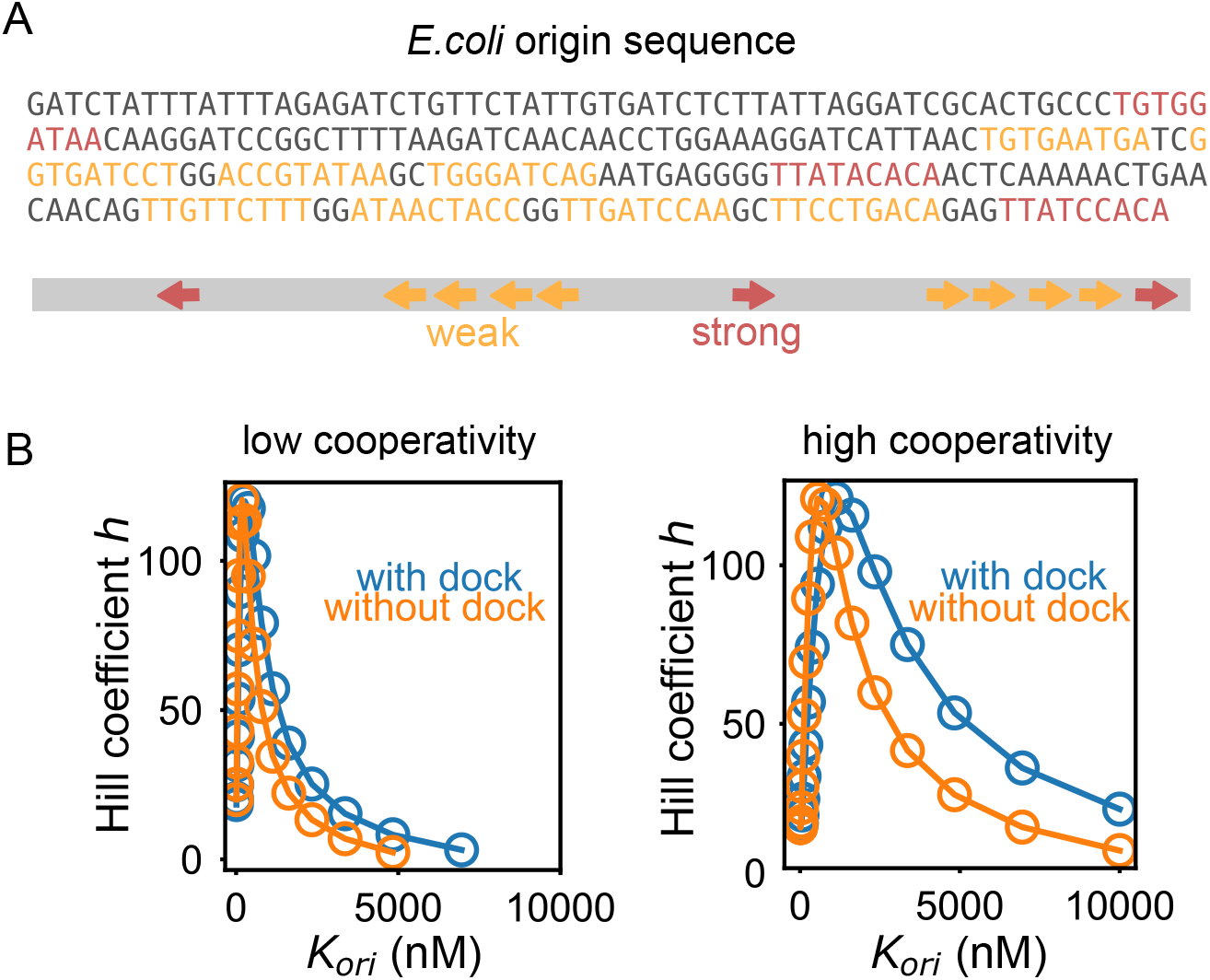
Docking sites expand the range of parameters that ensure high sensitivity. (A) Origin of replication in *E*.*coli*, with DnaA boxes represented in colors. The rightmost cluster is characterized by a sequence of weak DnaA boxes flanked by a strong one. (B) Hill coefficient as a function of the dissociation constant *K*_ori_ for low (*e*^*βJ*^ = 50) and high (*e*^*βJ*^ = 200) cooperativity. The two curves represent a case in which the origin is made of *m* = 8 weak binding sites (“without dock”), compared with a case in which the same number of weak binding sites are flanked by a strong docking site (“with dock”).

## Discussion

In this work, we identified the mechanisms underlying the ultrasensitive response in the DnaA system. In this perspective, chromosomal titration of DnaA is an instance of sequestration-based ultrasensitive response (***Wade Harper et al., 1993; Buchler and Louis, 2008; Buchler and Cross, 2009; Ferrell et al., 2014; Biswas et al., 2024; Sassi et al., 2020***). This mechanism leads to extremely large values of the Hill coefficient, provided that origin binding occurs right at the end of this titration process. The DnaA system is also characterized by cooperative interactions associated with a conformational change at the origin, reminiscent of a Monod-Wyman-Changeux mechanism, as well as direct interactions between DnaA-ATP molecules bound at neighboring sites. Although these cooperative mechanisms are classically associated with sharp response, they would not be able by themselves to lead to such high Hill coefficients in this system (***Owen and Horowitz, 2023***).

Our results contribute to understand why the DnaA system is regulated by two seemingly independent regulatory mechanisms (***Christensen et al., 1999; Kasho and Katayama, 2013; Katayama et al., 1998; Aranovich et al., 2006***). In particular, the main mechanism for cell cycle stability seems to be titration in slow growth and interconversion in fast growth (***Fu et al., 2023; Berger and ten Wolde, 2022, 2023; Knöppel et al., 2023; LaPorte et al., 1984***). Our results support this view, and show that these two different strategies naturally emerge by requiring ultrasensitive response in different growth conditions. Further, our model rationalizes why neither mechanism is effective in both slow and fast growth. Specifically, in fast growth, creation of chromosomal sites by preexisting forks limits the effectiveness of the titration mechanism at producing a sharp response. Additionally, the titration mechanism is prone to a dynamical instability in this regime (***Fu et al., 2023***). Conversely, our model predicts that the interconversion system is ineffective in slow growth, since its ability to produce a pulsatile response is based on sensing the concentration of replication forks already active before initiation. However, other mechanisms not explicitly included in our model that regulate the fraction of active DnaA (such as *datA*, DARS1, and DARS2) may play a role in slow growth condition (***Berger and ten Wolde, 2022***). More broadly, the framework introduced in this paper is flexible enough to incorporate other possible regulatory mechanisms, such as extrusion-regulated DnaA activity (***Li et al., 2025***).

Our model considers origin binding as the main source of stochasticity in the DnaA system. This choice is justified by the small number of origin binding sites compared to chromosomal ones. We expect that other sources of noise, for example fluctuations in total number of DnaA molecules, in number of molecules bound to chromosomal sites, and in the interconversion process, could affect the accuracy but without altering the qualitative picture presented here. Moreover, the nonuniform density of DnaA boxes along the chromosome may also play a role. A systematic analysis of these effects is left for a future study.

A key prediction of our theory is that ultrasensitivity can only be achieved in a narrow range of origin affinities. Testing this prediction would require measuring both the monomeric binding constant and the cooperativity of DnaA for the DnaA boxes at the origin. In *E. coli*, the dissociation constant for the strong sites has been estimated to be about 1 nM (***Schaper and Messer, 1995***), while the monomeric affinity for the weak sites has not been measured, to the best of our knowledge. Monomeric affinity of origin sites has been measured in *Streptomyces*, where such sites bind more strongly than in *E. coli* (***Majka et al., 2001***). The estimated values in *Streptomyces* are *βϵ* ≈ −5.3 and *βJ* ≈ −3.8 (***Majka et al., 2001***). These estimates are broadly consistent with the optimal predictions in Fig. 3a. Interestingly, *Streptomyces* DnaA binds with higher affinity to the strong *E*.*coli* boxes than to its own (***Majka et al., 2001***), supporting the view that the DnaA protein tends to be more conserved than the origin architecture (***Lu et al., 2023; Leonard and Mechali, 2013***). Quantitative measurements of binding constants in different organisms could provide a stringent test of our model predictions, and clarify the link between architecture of DnaA boxes and organism physiology.

## Methods

### Equilibrium binding

We consider a cell of volume *V* containing DnaA proteins at concentration *a*. We call *a*_atp_ and *a*_adp_ the concentrations of DnaA-ATP and DnaA-ADP, respectively, with *a*_atp_ + *a*_adp_ = *a*. A total of *n* = *V c*_tot_ chromosomal and *m* origin sites are available for DnaA binding. Using that *n* ≫ *m* and that *K*_ori_ ≫ *K*, we neglect the sequestering effect of the origin sites on chromosomal binding. Consequently, we first determine the equilibrium chemical balance between free and chromosomal-bound DnaA:

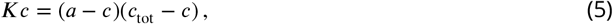

where *c* is the concentration of bound chromosomal sites. Since chromosomal binding sites have the same affinity for DnaA-ATP and DnaA-ADP, the concentration of chromosomal sites bound by DnaA-ATP is *c*_atp_ = *αc*.

The origin sites in the open and closed DNA conformation can be bound by DnaA, with dissociation constant *K*_ori_ and 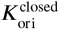, respectively. Monomeric binding is considered independent of the DnaA nucleotide state. Binding stabilizes the open conformation, in line with the Monod-Wyman-Changeux (MWC) mechanism of allosteric interaction. We assume that 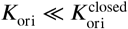.

Binding sites at the origin are found at short distance from each other. At these sites, the arginine finger of one protein (in either nucleotide state) can interact with the ATP of a neighboring DnaA-ATP protein, thereby strengthening binding (***Katayama et al., 2017***). We model this effect by a cooperative interaction characterized by a free energy *J*. Specifically, Dna-ATP can cooperatively bind to a bound protein (either DnaA-ATP or DnaA-ADP) immediately preceding it via arginine finger interaction (see Appendix). The ability to establish this cooperative interaction is the only model feature that differentiates DnaA-ATP from DnaA-ADP.

### Origin firing

An origin can fire when it is in the open conformation. The probability to be in the open conformation is expressed by

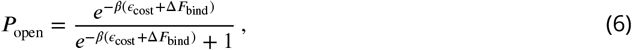

where *ϵ*_cost_ is the intrinsic free energy difference between the open and closed DNA conformation, and Δ*F*_bind_ is the free energy contribution due to preferential binding of DnaA in the open conformation. This free energy term is expressed by Δ*F*_bind_ = *k*_B_*T* ln *Z*^closed^/*Z*, where *Z* and *Z*^closed^ are the partition functions associated with the cluster of origin sites in the open and closed DNA conformations, respectively. We evaluated these partition functions using the transfer-matrix method (see Appendix and (***Hill, 1985***)). Such method allows for explicitly computing the partition function for different origin architectures, including the case of multiple clusters and with the presence of docking sites. Unless specified otherwise, results in the Main Text are for the case of a cluster of *m* weak sites, for which we obtain

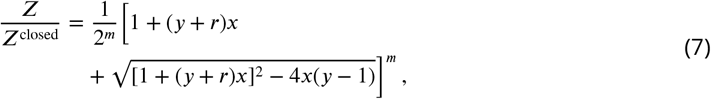

where *x* = *α*(*a* − *c*)/*K*_ori_, *y* = *e*^*βJ*^ 290, and *r* = (1 − *α*) / *α* The stochastic rate of initiation is given by *k* = *P*_open_*k*_max_, where *k*_max_ is a rate constant.

### Fraction of active DnaA

Multiple regulatory processes modulate the fraction of active DnaA-ATP over the cell cycle (***Knöppel et al***., 2023; ***Boesen et al., 2024***). In particular, the fraction of DnaA-ATP peaks around the time of initiation, and drops immediately after due to ATP hydrolysis, promoted by a replication fork-dependent mechanism known as regulatory inactivation of DnaA (RIDA) (***Knöppel et al., 2023; Suetsugu et al., 2005***). Even in slow growth conditions, where replication forks are absent during a substantial fraction of the cell cycle, additional DNA-dependent hydrolysis pathways keep the active fraction of DnaA relatively low (***Fu et al., 2023***). These effects have been modeled in detail in previous work (***Berger and ten Wolde, 2023***). We take into account these effects via a simple dynamics of the form

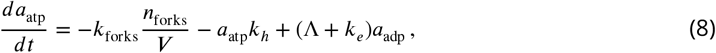

where *a*_adp_ is the total concentration of DnaA-ADP, *n*_forks_ is the number of active forks, *k*_forks_ is the forkmediated hydrolysis rate constant, *k*_*h*_ is the intrinsic hydrolysis rate, *k*_*e*_ is the exchange rate, and the term proportional to the growth rate Λ accounts for dilution. Assuming that the dynamics is close to a steady state and that fork-mediated hydrolysis operates close to saturation, we approximate the solution of Eq. (8) by

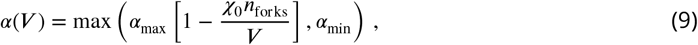

where *α*_min_ and *α*_max_ are two nutrient-dependent limits, and χ_0_ = *k*_forks_/[*a*(Λ + *k*_*e*_)] quantifies the relative strength of fork-coupled hydrolysis compared to the rate of nucleotide exchange (ADP → ATP), see Appendix.

### Cell cycle dynamics

In the model, we take the replication time *C* and the time *D* between termination and division as constant (***Cooper and Helmstetter, 1968***). We assume that the chromosomal sites are uniformly distributed along the genome (***Fu et al., 2023; Berger and ten Wolde, 2023***). The cell volume between divisions grows exponentially: *V* (*t*) = *V*_0_ exp(Λ*t*), where *V*_0_ is the volume at birth, *t* is the time since birth and Λ is the growth rate. The total number of titration sites for a given number of forks *n*_forks_ increases at a rate *n*_forks_*g* / (2*C*), where *g* is the total number of sites in one chromosome. At every instant, each of the eligible origins can independently fire at rate *k*. Replication begins at the selected origin after a short delay *t*_delay_, and the origin subsequently enters an eclipse period *t*_eclipse_ during which reinitiation is blocked (***Campbell and Kleckner, 1990; Lu et al., 1994***). Upon division, the volume is halved, and one of the genomes is randomly selected for the tracked daughter cell.

### Model parameters

Numerical values of the parameters are reported in Table 1.

### Hill coefficient

Combining Eq. (1) and Eq. (6), we express the Hill coefficient by

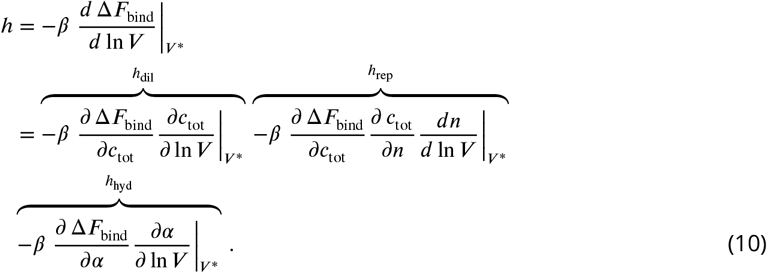

Using Eq. (1) and the expression for Δ*F*_bind_ in terms of the partition function (Eq. (7)) we obtain, after some algebra, that the initiation volume is expressed by

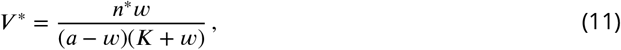

see Appendix. We then evaluate the different contribution to the Hill coefficient in Eq. (10) using the chain rule (see Appendix). For example, *h*_dil_ is computed by expressing ∂ Δ*F*_bind_/∂*c*_tot_ = (∂ Δ*F*bind/∂*c*)(∂*c*/∂*c*_tot_ ), and using Eq. (7) and Eq. (5) to evaluate the two partial derivatives (see Appendix). This procedure results in the expression for *h*_dil_ given in Eq. (4). For *h*_rep_, we use that *dn*/*dt* = *n*_forks_*vp*, where *ρ* is the linear density of chromosomal binding sites, and that 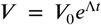. Applying the chain rule again we find

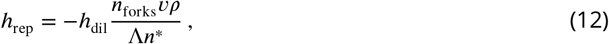

see Appendix. Finally, using Eq. (9) into Eq. (10) we obtain

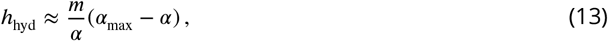

see Appendix.

### Special sites

We consider the case in which a fraction *θ* of chromosomal sites can be only bound by DnaA-ATP. In this case, Eq. (5) is generalized into

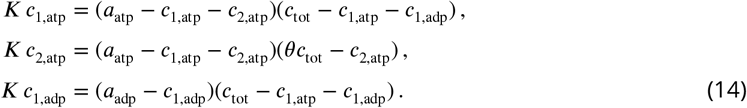

where the indices 1 and 2 denote ordinary and special sites, respectively. Results presented in the Main Text are obtained by solving Eq. (14) for small *θ* using a perturbative expansion (see Appendix).

## Author contributions

A.S. and S.P. designed and performed the research and wrote the paper. A.S. wrote the code and performed numerical simulations.

## Acknowledgments

We are grateful to Ariel Amir, Kuheli Biswas, Marco Cosentino Lagomarsino, Mattia Corigliano, Suck-joon Jun, Shogo Ozaki, Samuel Hauf, Bianca Sclavi, David Fange, Florian Pflug, Irene Pellini, Kazutoshi Kasho and Yuhai Tu for insightful discussions. We thank Pieter Rein ten Wolde and Haochen Fu for precious feedback on a preliminary version of the manuscript.

## Data availability

Mathematical derivations of the results of the paper are available in Appendix. Code to simulate the model is available at https://github.com/alberto-sassi/Ultrasensitivity_in_DnaA_system.

## Appendix Partition function and transfer matrix

### Probability of the open conformation

The firing rate *k* is expressed by *k* = *k*_max_*P*_open_. At thermodynamic equilibrium, the probability of finding the DNA at the origin in the open conformation is given by

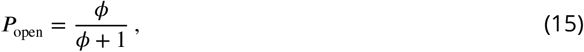

where

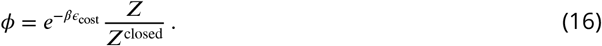

For appropriate parameters, *P*_open_ presents a sharp transition from *P*_open_ ≈ 0 to *P*_open_ ≈ 1 as the volume is increased. We approximate this dependence with a Hill function

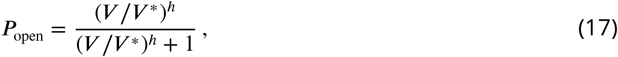

where *h* is the Hill coefficient. Comparing Eq. (16) and Eq. (17), we obtain

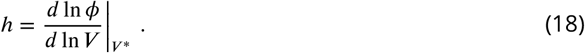

We shall derive an explicit expression for Φ by evaluating the partition functions. We then will use the result to compute *h* using Eq. (18).

Computation of the partition function using the transfer matrix method

We here compute the partition functions *Z* and *Z*^closed^ for different boundary conditions using the transfer matrix method. The transfer matrices for our model are

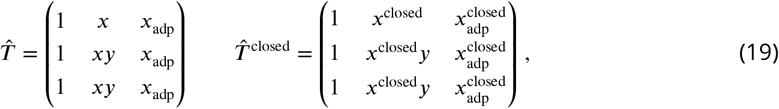

where

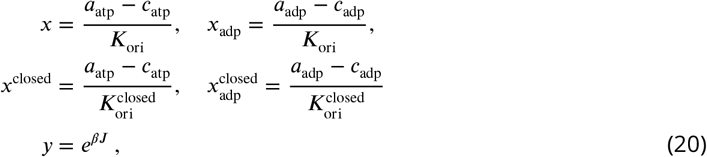

where *K*_ori_ and 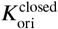 are the dissociation constant at the weak origin sites in the open and closed DNA conformation, respectively. Since the two transfer matrices have the same structure, we focus for simplicity on the one for the open conformation.

The two non-vanishing eigenvalues of the transfer matrix are

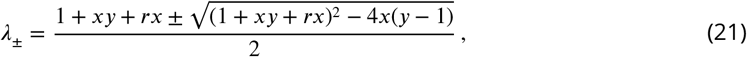

where *r* = *x*_adp_/*x*. Both eigenvalues are increasing functions of *x*, and we have *λ*_−_ < 1 < *λ*_+_.

The transfer matrix is diagonalized by a transformation 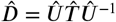, with

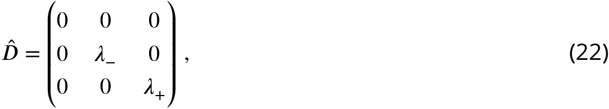

and

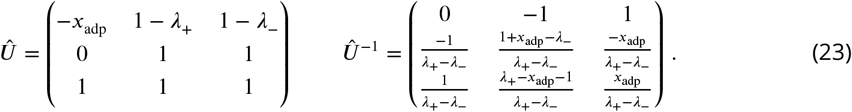

We now consider three specific scenario, corresponding to different boundary conditions.

**Appendix 0—figure 6.**
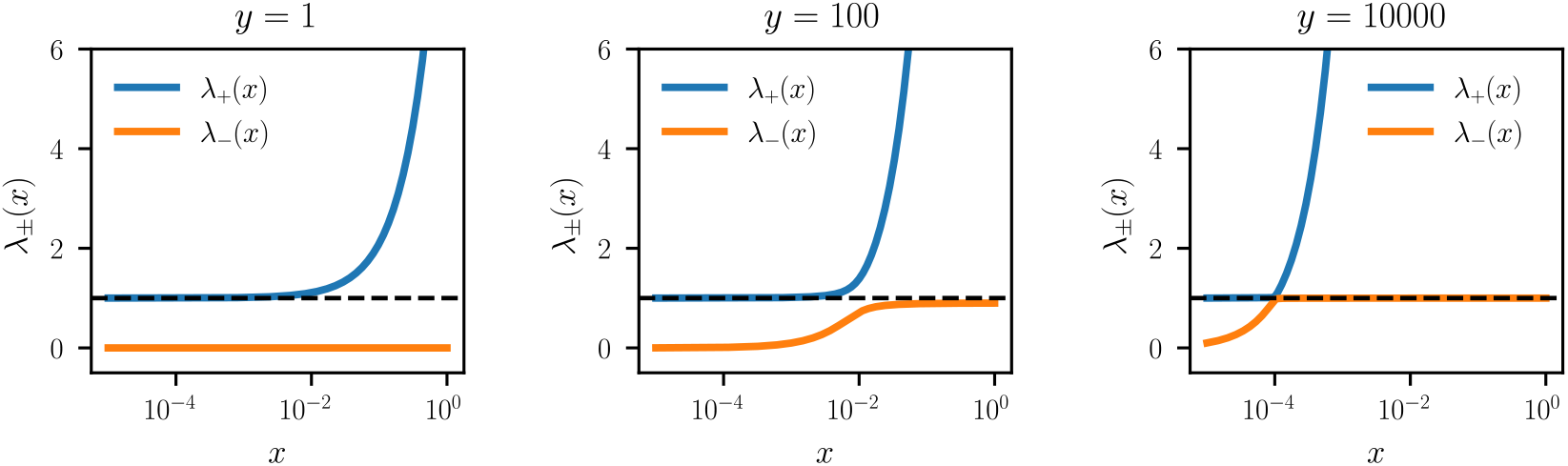
Eigenvalues of the transfer matrix as a function of *x* for different values of the cooperativity weight *y* assuming *r* = 0.

### Periodic cluster (ring) of weak sites

We consider a ring of *m* weak sites. In this case, we have

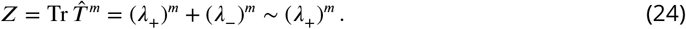

The approximation comes from the observation that *λ*_+_ is on the order of *x*(*y* + *r*), while *λ*_−_ is on the order of *y*/[*x*(*y* + *r*)^2^]. This result implies that

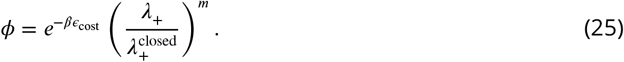

We now discuss whether neglecting the contribution from the second eigenvalue is appropriate. In the case of strong cooperativity, the two eigenvalues are close to each other for a certain value of *x*, see Fig. 6). In contrast, in the absence of cooperativity (*y* = 1), *λ*_−_ = 0 and *λ*_+_ = 1 + *x*.

#### Linear cluster of weak sites

The origin is not a ring but a linear sequence of binding sites. For the sake of simplicity, we assume that the origin is formed by a single linear cluster of *m* sites. Extension to a more detailed architecture that includes several smaller clusters relatively far apart would not significantly change the main results of this work. We consider a single linear cluster of *m* sites. The partition function is given by

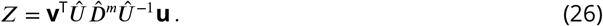

With

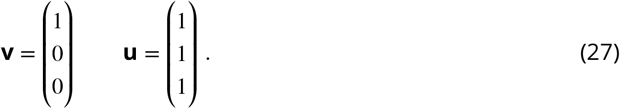

The first matrix multiplications are

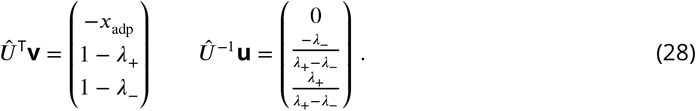

We plug these expressions into Eq. (26), obtaining

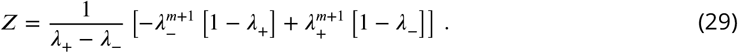

#### Linear cluster of weak sites flanked by a strong docking site

We consider a linear cluster having a strong site to its side. In this case, *m* represents the number of weak sites in the cluster. The partition function is still given by Eq. (26), but the boundary vectors **v** and **u** need to be modified:

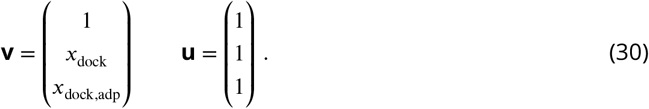

Here, *x*_dock_ = (*a*_atp_ −*c*_atp_)/*K*_dock_ and *x*_dock,adp_ = (*a*_adp_ −*c*_adp_)/*K*_dock_, where *K*_dock_ is the dissociation constant associated with the docking site. We assume that the dissociation constant for the docking site is the same in the two conformations, so that we can measure specifically how the docking site stabilizes the subsequent occupation of the adjacent weaker sites.

We now compute

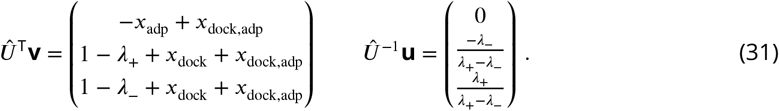

We plug these expressions into Eq. (26), obtaining

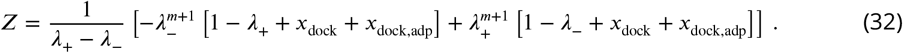

When there is no cooperativity (*y* = 1), we have *λ*_−_ = 0 and the correction of the docking sites provides a similar prefactor in the term for the open and the closed conformation:

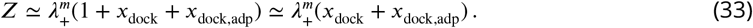

In the presence of cooperativity, all terms in the right hand side of Eq. (32) contribute in the equation for the open conformation.

Conversely, for the closed conformation we can still use the approximation *x*^closed^ ≪ 1 and thus

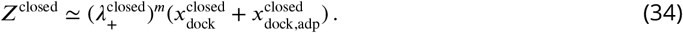

#### Calculation of the Hill coefficient

##### Calculation of the threshold concentration

We calculate the midpoint volume *V* ^*^ and 0074hen use Eq. (18) to find the Hill coefficient *h*. We start by finding the midpoint value of *x, x* = *x*^*^, see Eq. (20).

We define

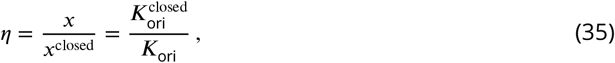

so that Eq. (16) becomes

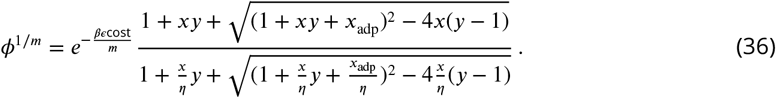

We assume that η is large, so that the expression reduces to

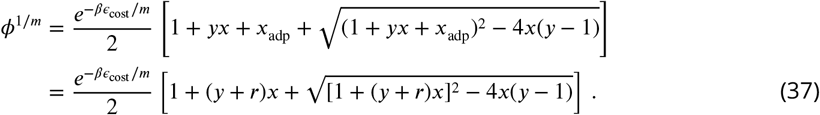

We expect to have more free DnaA-ATP than DnaA-ADP at initiation, so that *r* should be less than 1 for *V* = *V* ^*^. We also define 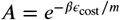 for convenience of notation.

We now impose Φ = 1 for *x* = *x*^*^, obtaining

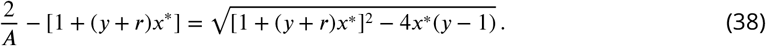

Squaring both sides of the equation we get

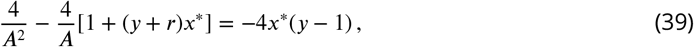

which simplifies to

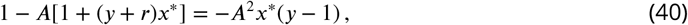

so that

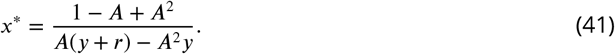

We now assume that *y* ≫ *r* and, since *A* < 1, we neglect the *A*^2^ in the numerator, obtaining

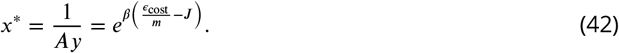

This calculation shows that, under our assumptions, the value *x* = *x*^*^ is unaffected by the presence of DnaA-ADP, because the concentration of DnaA-ADP is not large enough to initiate replication without cooperative binding.

Equilibrium occupancy of the chromosomal sites

The concentrations of occupied sites are given by detailed balance:

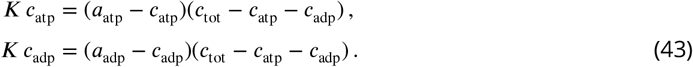

Summing the two equations in Eq. (43), we obtain

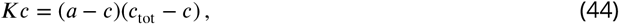

where *c* = *c*_atp_ + *c*_adp_ . The solution of this second order algebraic equation is

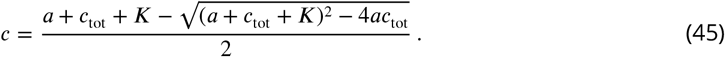

The proportion of sites bound to DnaA-ATP and DnaA-ADP are

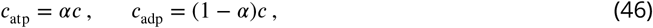

where *α* = *a*_atp_/*a*. Indeed, looking for solutions of the form *c*_atp_ = ξ*c* in the first equation, we obtain

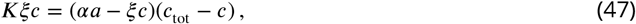

and we need *α* = ξ for Eq. (47) to be compatible with Eq. (44). Equation (46) is a consequence of the fact that the dissociation constant for the sites is assumed to be the same for the two nucleotide states of DnaA, therefore the partitioning of the proteins bound to the chromosomal sites matches the total proportions of proteins in the cell.

Volume at midpoint at fixed fraction of active DnaA

We now find an explicit expression for *V* ^*^ at fixed *α*. We define

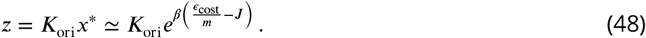

Given the definition of *x, z* is equal to the concentration of free DnaA-ATP at initiation:

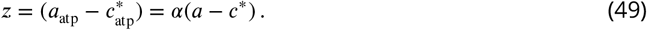

Combining Eq. (48) with Eq. (49) we obtain the condition for the midpoint

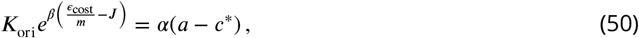

where the left hand side depends on the origin parameters, while the right hand side contains the effect of titration.

Solving Eq. (45) in 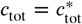 and using Eq. (49) to eliminate *c*^*^ we obtain

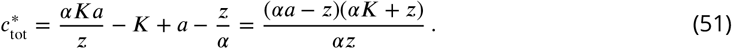

Therefore, the volume at the midpoint is expressed by

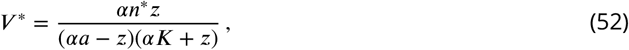

where we have defined the number of titration sites at initiation 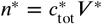.

Calculation of the Hill coefficient at fixed fraction of active DnaA and number of titration sites

We calculate the Hill coefficient in case *h*_rep_ = *h*_hyd_ = 0, using Eq. (18) and the chain rule:

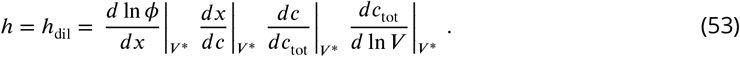

We calculate the derivatives appearing in the right hand side of Eq. (53) one by one. At the moment we keep the active fraction *α* and the total number of titration sites *n* fixed. The first term in Eq. (53) is given by

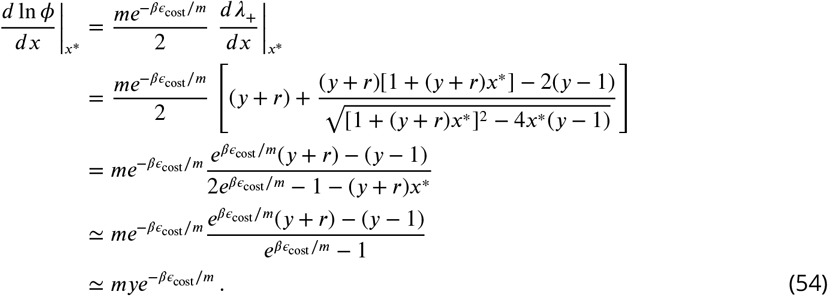

In the third line, we have used that 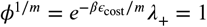 at the midpoint, so that

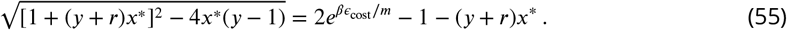

We now move to the second term. From the definition of *x* in Eq. (20) and Eq. (46) we immediately obtain

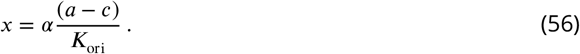

We therefore have

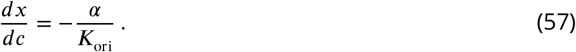

For the third term, using Eq. (45) we have

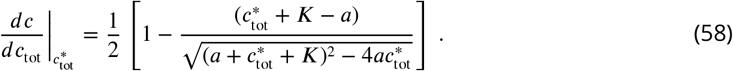

To simplify the square root term, we use again Eq. (45):

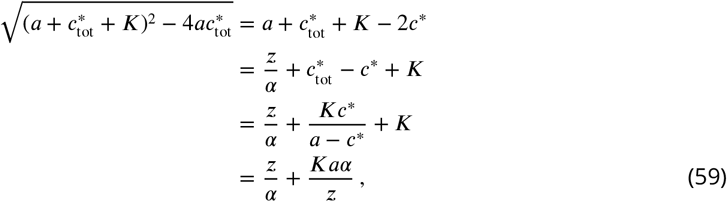

where we have used Eqs. (44) and (49). Substituting this result into Eq. (58) we obtain

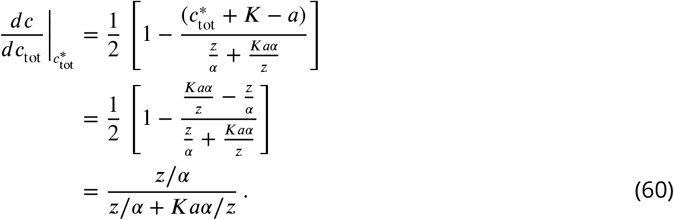

Finally, since *c*_tot_ = *n*/*V*, we have

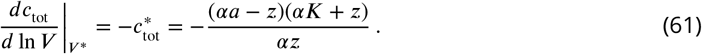

Substituting all of these results into Eq. (53), we have

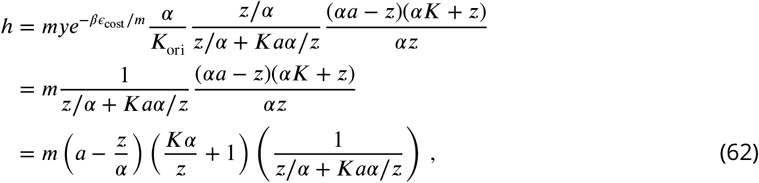

and, defining the ratio *w* = *z*/*α*:

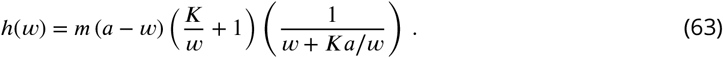

Optimal Hill coefficient at fixed fraction of active DnaA and number of titration sites We look for the value of *w* that maximizes the Hill coefficient:

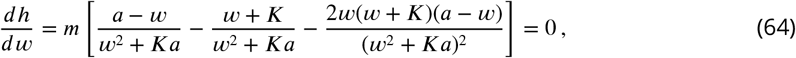

obtaining

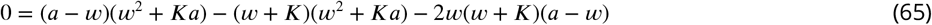

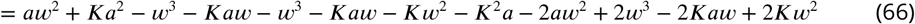

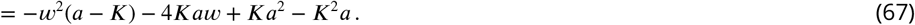

Using that *a* ≫ *K*, we obtain

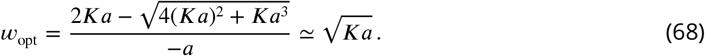

This is also the value of *w* for which the total concentration of initiator proteins is equal to the concentration of binding sites. Indeed, if we consider Eq. (51) with the condition

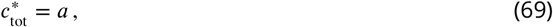

we obtain

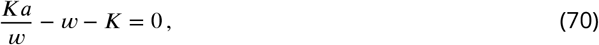

and thus

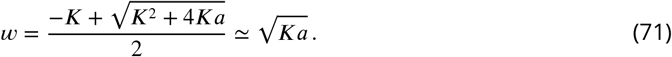

Substituting this value back into Eq. (63) we estimate the optimal value of the Hill coefficient

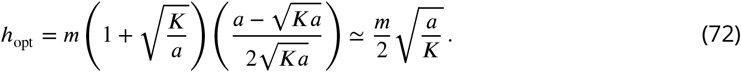

Hill coefficient for varying fraction of active DnaA and number of titration sites In general, the Hill coefficient can be decomposed as

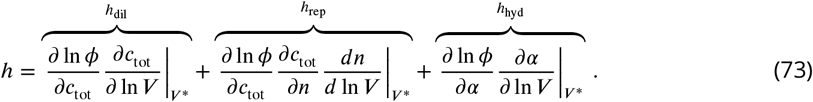

We have already derived *h*_dil_. We now derive the other two terms.

Calculation of the Hill coefficient associated with changing fraction of active DnaA In the model, the fraction *α* is determined by

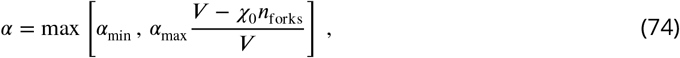

where *n*_forks_ is the number of replication forks, and *α*_min_, *α*_max_ and χ_0_ are positive constants. We have

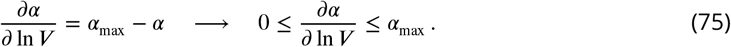

In slow growth, before initiation one has *n*_forks_ = 0, so that *α* = *α*_max_ and ∂*α*/∂ ln *V* = 0.

We now return to the general case. Since *r* = (1 − *α*)/*α*, we have

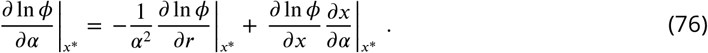

We start by the first term:

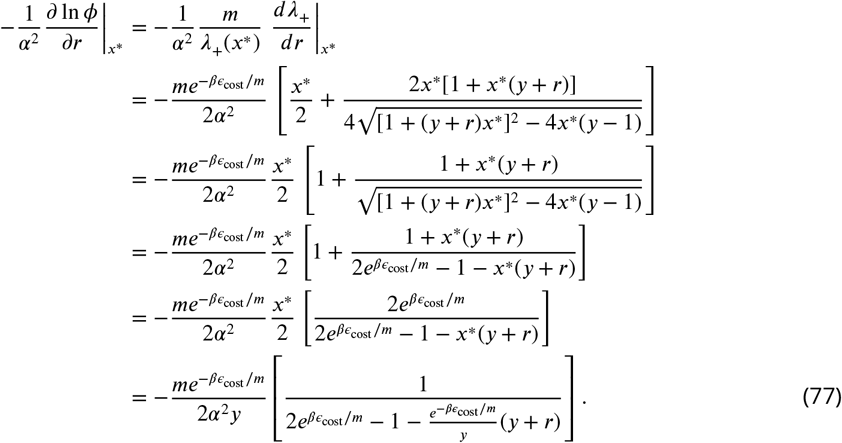

We now move to the second term:

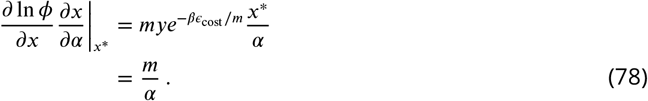

We now use these results to empirically estimate *h*_hyd_ for *E. coli* in optimal growth. Realistic parameters are *y* ≈ 50, *βϵ*_cost_ ∈ (10, 20), *m* ≈ 8, *α* ≈ 0.15, which give

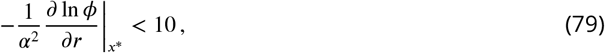

and

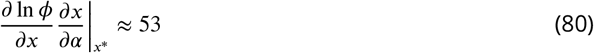

so that the second term dominates over the first. We also have ∂*α*/∂ ln *V* ≈ 0.85, so that

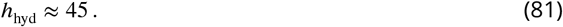

Contribution to the Hill coefficient associated with changing number of titration sites The number of chromosomal sites varies as

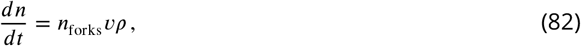

where *n*_forks_ is the number of active forks, *v* is their speed, and *ρ* is the density of chromosomal sites along the genome. Given that *c*_tot_ = *n*/*V*, we have

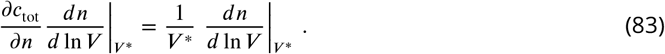

Using the dependence of the volume on the growth rate, Λ*dt* = *d* ln *V*, we simply have

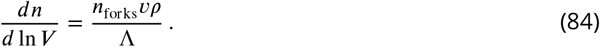

Using the fact that

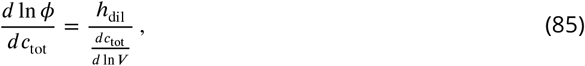

we have

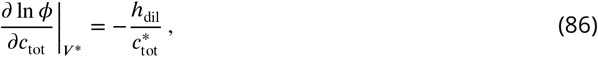

and thus

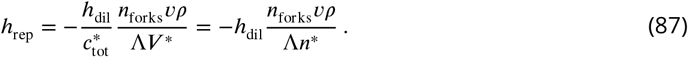

We estimate *h*_rep_ for E.coli in optimal growth conditions (τ = 25 min). We first determine whether initiation takes place before or after termination. The time of initiation in the regime *τ* < *C* < 2τ is given by

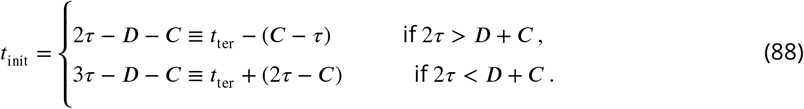

We assume *D* = 20 min, so that 2τ = 50 < *D* + *C*. Therefore, the time of termination is *t*_ter_ = *τ* − *D* = 5 min and the time of initiation is *t*_init_ = 3τ − *D* − *C* = 15 min. There are two genomes in the cell at the time of initiation. The replication progress between two initiations is τ/*C* in steady state. Therefore, if the total number of sites per chromosome is *n*_sites_ = 300, then

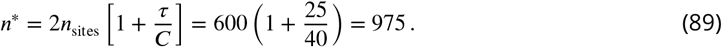

Before initiation, we have

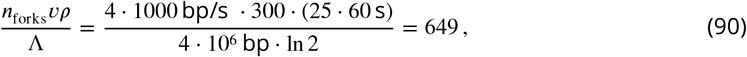

and thus

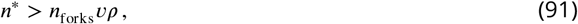

right before initiation. Then, after the initiation, the number of forks moves from *n*_forks_ = 4 to *n*_forks_ = 12 and

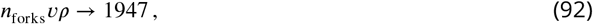

and thus

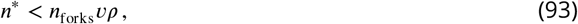

right after initiation.

Now we make an estimate of *h*_rep_ in these conditions, under the assumptions made for Eq.(81).

Before initiation, we have

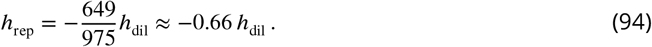

Therefore, if *h*_dil_ = 100, the value of the total Hill coefficient in fast growth becomes

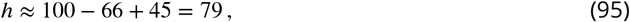

where we have used Eq. (81) for the contribution *h*_hyd_.

#### Stability analysis

##### Midpoint volume at varying fraction of active DnaA

We derive a formula for *V* ^*^ that, in contrast with Eq. (52), explicitly takes into account that *α* depends itself on *V*, see Eq. (74). Close to initiation, when *α* is growing, we have

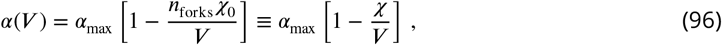

where we defined χ = *n*_forks_χ_0_. The expression for the midpoint volume becomes

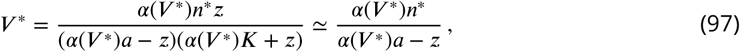

where the approximation follows from the fact that *K* is small. Solving for *V* ^*^, and keeping only the biologically admissible solution for which *α* > 0, we obtain

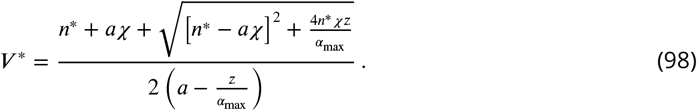

We now use the facts that *V* ^*^ is close to *n*^*^/*a* at initiation and that *α* is a small number. Therefore, we expand *n*^*^ around *n*^*^ = *a*χ to obtain

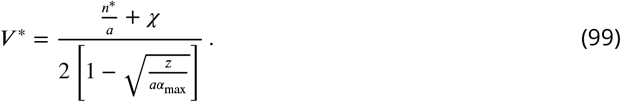

We note that the affinity for the origin binding sites should always be such that 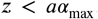 Since *z* is the concentration of free DnaA-ATP at the midpoint, this constraint means that the maximal concentration of DnaA-ATP must be large enough to trigger initiation.

##### Conditions of stability in case of multi-fork replication

A sharp response is not enough to ensure proper timing of replication. In case of multi-fork replication, small deviations from the proper time interval between initiation events might be amplified from generation to generation even when the firing rate is a step function of the volume. In this section we perform a stability analysis assuming stepwise response at *V* ^*^, to determine the conditions on the RIDA mechanism to ensure precise timing of initiation. A similar analysis in the absence of hydrolysis was first proposed by Fu et al. (***Fu et al., 2023***) to show that the system is not stable in the regime *τ* < *C* < 2τ, where *τ* is the doubling time and *C* the replication time.

Initiation is triggered whenever

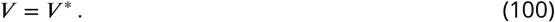

We need to find the relation between two subsequent intervals between consecutive initiations. Let *t*_*a*_ be the time between two initiations and *t*_*b*_ the time to be waited for a third one afterward. By construction, we have

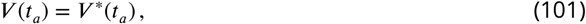

and

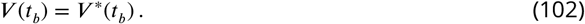

Moreover, we also use the fact that *V* increases in time exponentially at rate Λ, whereas *V* ^*^ evolves according to Eq. (99). In the regime *τ* < *C* < 2τ, for every genome the number of sites right before the first initiation is given by the sum of the sites of one full chromosome and the newly created sites during the time *t*_*a*_:

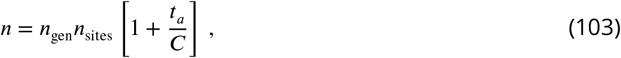

where *n*_sites_ is the number of sites of one full chromosome and *n*_gen_ is the number of genomes in the cell right before initiation. We have

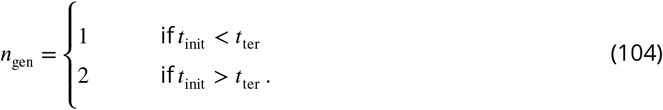

**Appendix 0—figure 7.**
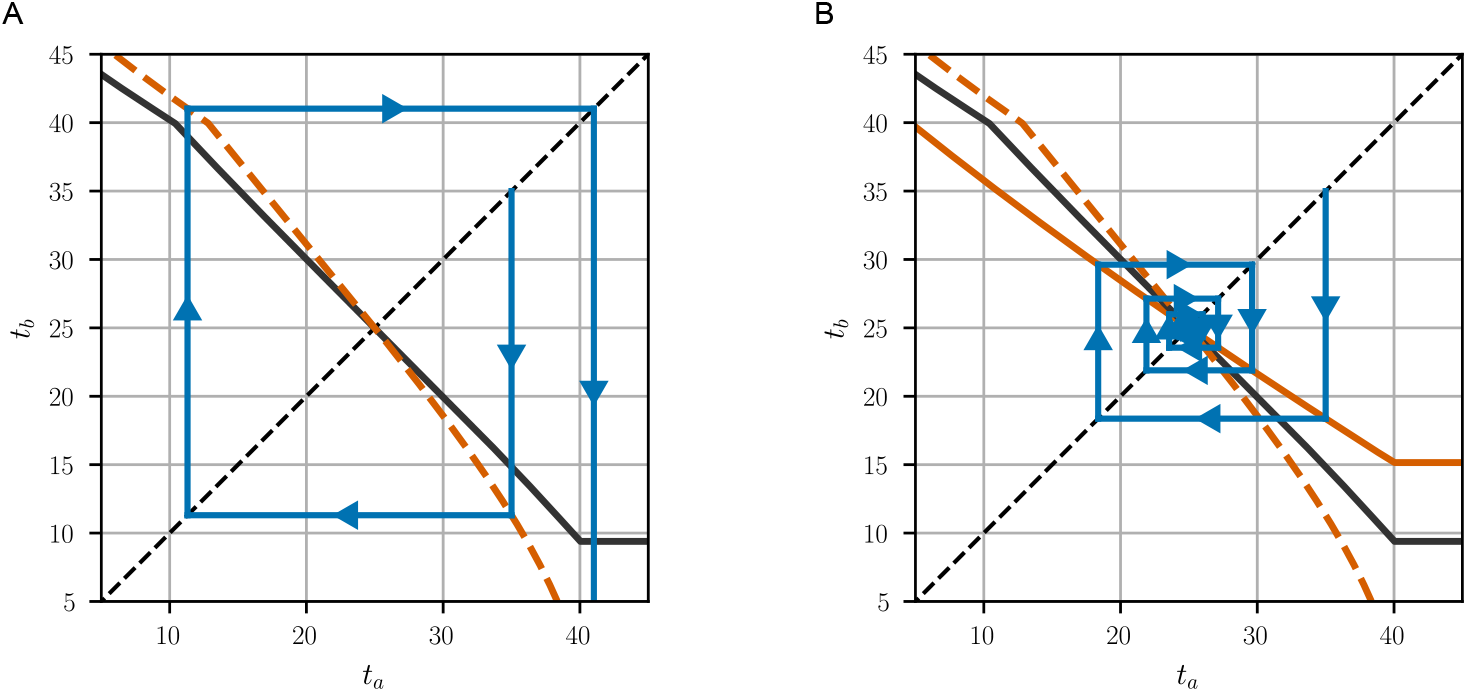
Cobweb plot for consecutive intervals between initiation events in case 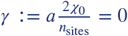 (A) and γ = 0.6 (B). The orange lines are given by Eq.(106) in case γ = 0 (dashed) and γ = 0.6 (continuous). The black continuous line was obtained using the threshold value for χ_0_, expressed in Eq.(108).

The total number of forks in the regime *τ* < *C* < 2τ is

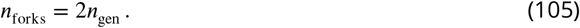

We can now use Eq. (101) and Eq. (102) to find the relation between two consecutive intervals between initiation events:

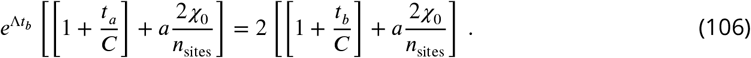

The stability condition is

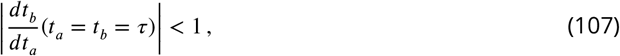

resulting in

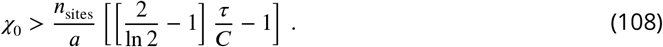

Only when Eq. (108) is satisfied the intervals between initiations converge (Fig.7).

#### Special chromosomal sites

In this section, we consider a variant of the model in which DnaA-ATP has access to sites that are not accessible to DnaA-ADP, but with the same dissociation constant *K*. We use the index 1 to indicate the ordinary sites that can be occupied by DnaA in either state and the index 2 for the other special sites. We call *c*_tot_ the total concentrations of the ordinary sites and

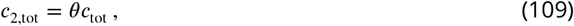

the total concentration of special sites, where *θ* is a constant.

The chemical balance reads

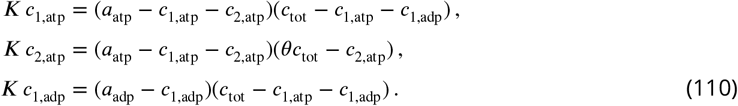

where the index for the concentration of occupied binding sites represents the nucleotide state of the protein bound to the given binding site, therefore, for example, *c*_1,adp_ is the concentration of ordinary sites bound by DnaA-ADP. We assume that the special sites are few compared to the ordinary sites, so that we can look for solutions for small θ:

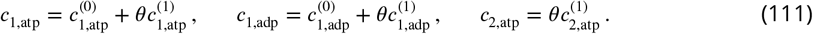

The solutions are found by plugging these equations into Eqs. (110) and by matching orders in θ.

For the zeroth order 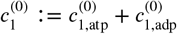 we use the result we obtained in the previous sections:

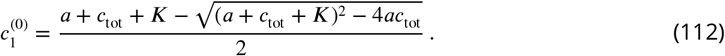

Then, we simply have 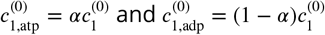.

We now proceed with the first order terms. We start with the occupation of special sites *c*_2,atp_.

The equation for the first order in *θ* is

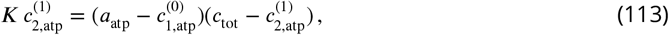

and the solution is

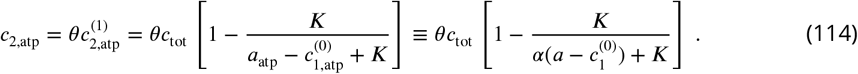

As expected, the concentration of occupied special sites is closer to their total number the smaller is *K* compared to the concentration of free proteins. Before we find 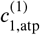 and 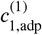, it is useful to obtain their sum:

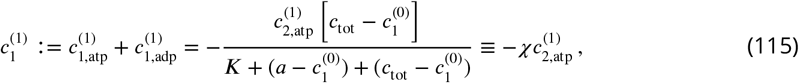

where we have defined χ to separate 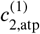 from the rest of the expression. Note that 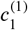 is always negative, because the addition of the special sites reduces the available pool of proteins. The equations for the remaining concentrations 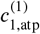 and 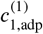 can be solved in terms of 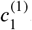. In the case of DnaA-ADP we have

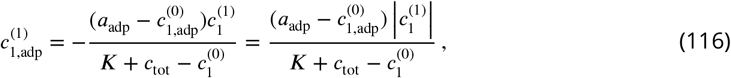

which is always positive because, thanks to the special sites titrating more DnaA-ATP, the DnaA-ADP proteins have less competition for the ordinary sites, and

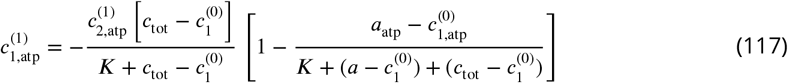

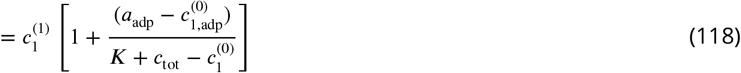

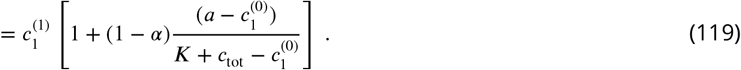

The total concentration of sites occupied by DnaA-ATP is

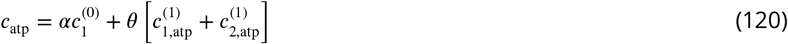

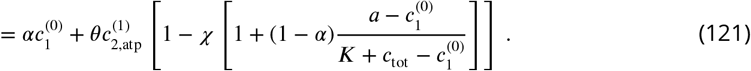

Writing explicitly χ and 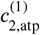, we obtain

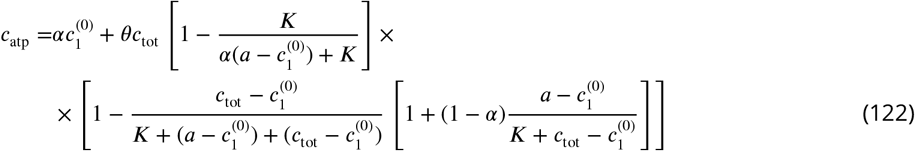

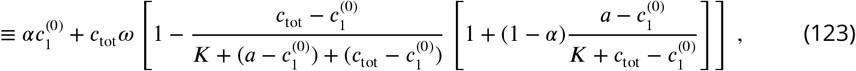

With

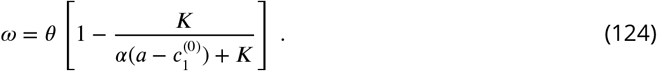

We use the fact that

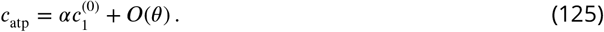

Thanks to Eq. (125), when considering the value at the midpoint, the approximation

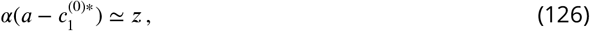

holds for every term contained in the expression multiplying θ, because any correction would be of the order of θ^2^. Additionally, we have

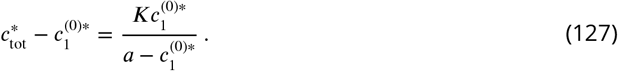

Using these expressions, we rearrange the formula:

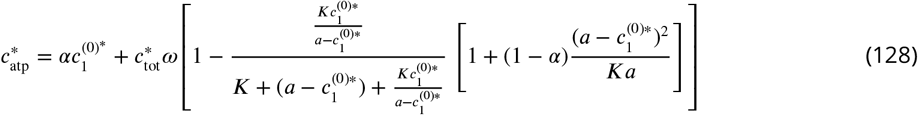

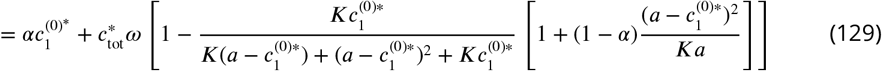

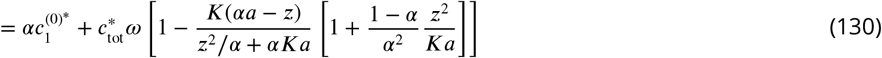

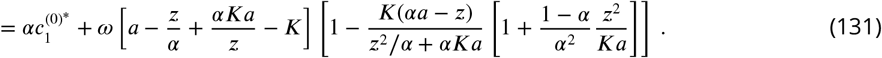

To solve this equation, we are left with the only term containing 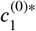 for which we cannot use Eq. (126), and we rather have to use the solution of the mass balance equation for *c*, namely Eq. (45). First, we lighten the notation by defining

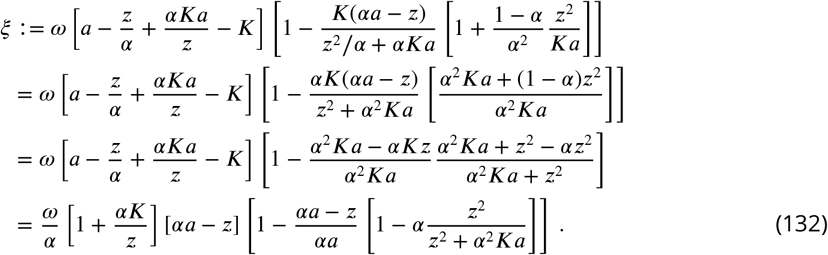

We use this definition of ξ to obtain an expression for 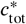 :

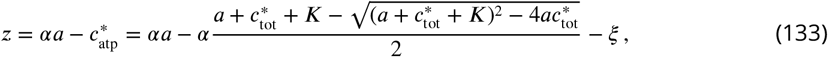

and thus

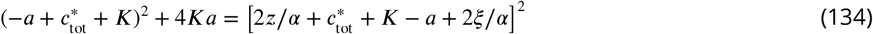

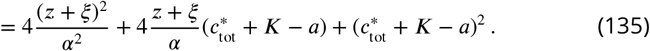

By solving for 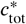 we have

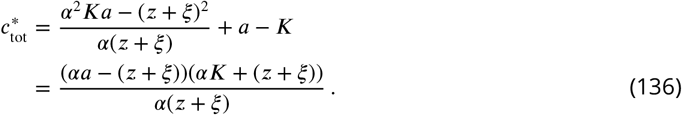

Note that when ξ = 0 we recover the formula obtained previously for *c*^*^ .

The volume at initiation is

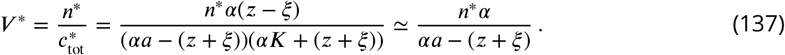

We expand for small ξ to obtain

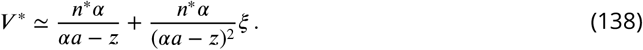

The contribution from the special binding sites is

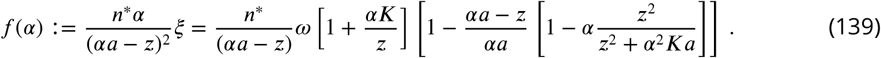

The function in Eq. (139) is a monotonically decreasing function of *α*, showing that the effect of deactivation is enhanced by the special binding sites.

#### Derivation of the fraction of active DnaA from a mechanistic model

The process by which DnaA is activated and de-activated through reactions of ATP hydrolysis and nucleotide exchange has been studied extensively. The known regulatory mechanisms are RIDA and datA for the hydrolysis of ATP, and DARS1/DARS2 and the membrane phospholipids for the exchange from ADP to ATP. However, it was shown (***Boesen et al., 2024; Knöppel et al., 2023; Kasho et al., 2017***) that the system can function even without datA and DARS1/DARS2, that play a supporting role. In this section we derive the fraction of acive DnaA under the simplifying assumption that RIDA is the only active regulatory mechanism involved in the interconversion between ATP and ADP. Additionally, a DnaA-ATP molecule can spontaneously convert to DnaA-ADP at a lower rate. The main factor restoring the active state is assumed to be the synthesis of new DnaA proteins, that are more likely to bind ATP than ADP, because ATP is more abundant.

The main characteristic of RIDA is that the hydrolysis is related with the presence of the replication fork. Consequently, previous models (***Fu et al., 2023; Berger and ten Wolde, 2022***) incorporate hydrolysis via Michaelis Menten equation that depends the number of replication forks *n*_forks_:

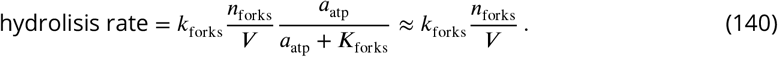

where *k*_forks_ and *K*_forks_ are constants, and the approximate equality descends from the assumption that RIDA operates near saturation. The corresponding dynamical equation for the concentration of active DnaA is

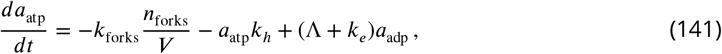

**Appendix 0—figure 8.**
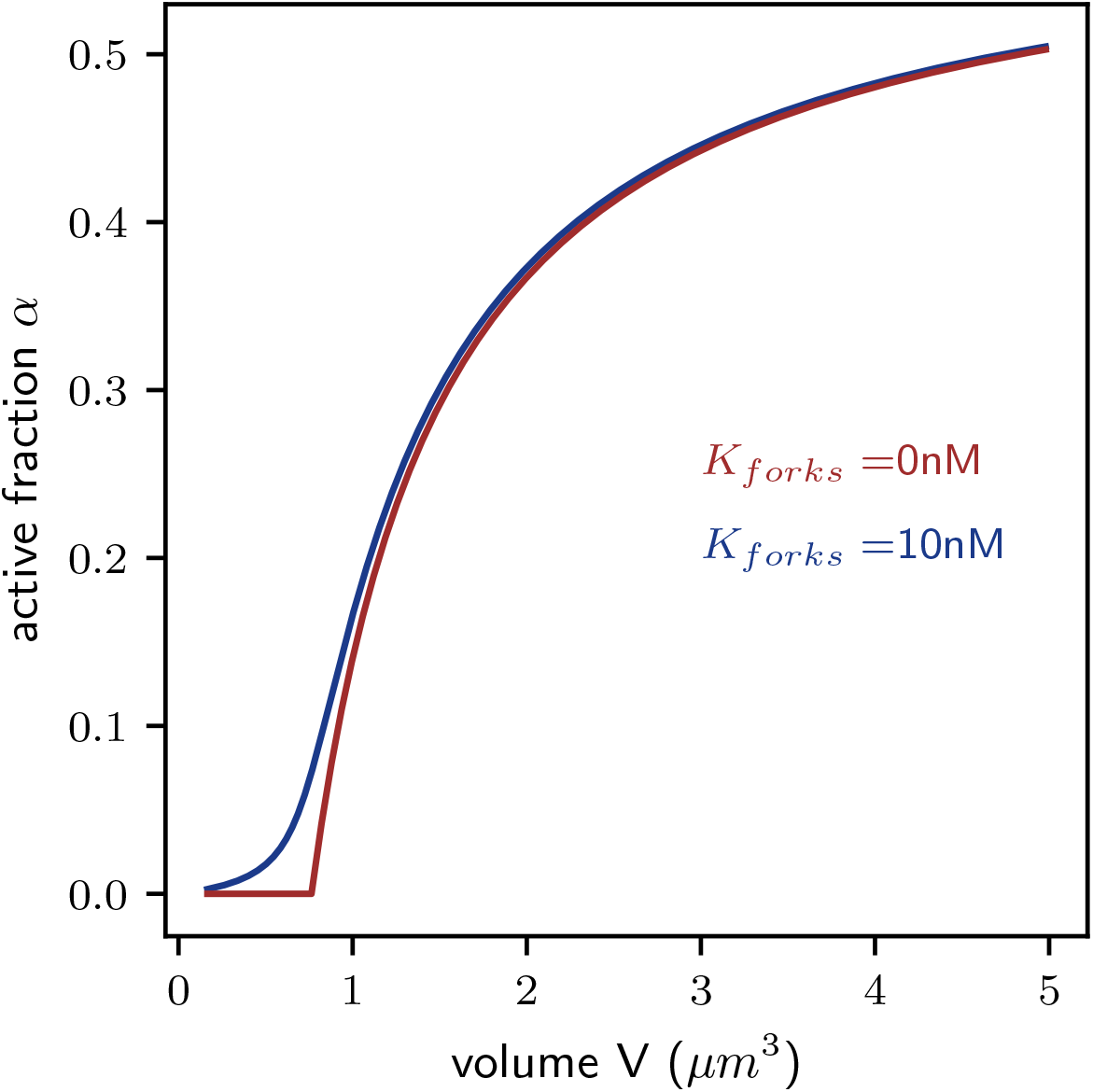
Comparison of the steady state solution of the equation for *α* in the case a Michaelis Menten kinetics (blue) or a zeroth order reaction kinetics (red) are used.

where *k*_*h*_ and *k*_*e*_ are the intrinsic rates of hydrolysis and synthesis, respectively, and Λ is the growth rate. The equation is coupled with the conservation law *a*_atp_ + *a*_adp_ = *a*. At steady state, we have

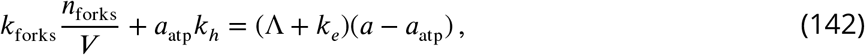

and thus

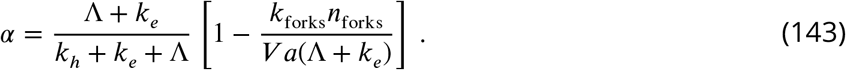

Defining

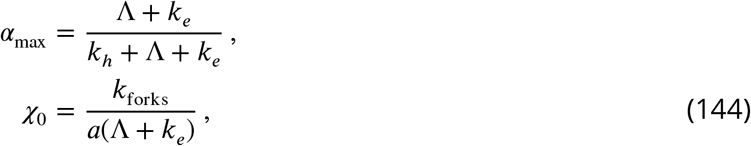

we obtain the phenomenological relation in the Main Text for *α* > *α*_min_:

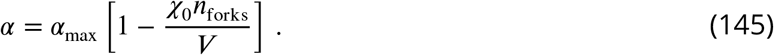

Considering a zero-th order reaction for the hydrolysis (*a*_atp_ ≫ *K*_forks_) is no longer possible as *α* approaches zero. A comparison with the curve without approximation (Fig. 8) shows that the deviation is only relevant far from the volume at initiation in fast growth conditions in case *K*_forks_ = 10 *nM*(> 1*μm*^3^, see ***Wallden et al. (2016***)).

