## Supplemental material for "Ultrasensitive response in bacterial replication initiation" (from another version of this preprint: version 2, 2026-09-15)

### Supplemental material for manuscript "Ultrasensitive response in bacterial replication initiation"

**4 Special chromosomal sites** **11**

**5 Derivation of the fraction of active DnaA from a mechanistic model** **14**

### 1 Partition function and transfer matrix

#### 1.1 Probability of the open conformation

The firing rate $k$ is expressed by $k = k_{\max} P_{\rm open}$. At thermodynamic equilibrium, the probability of finding the DNA at the origin in the open conformation is given by

$$P_{\rm open} = \frac{\phi}{\phi+1}\,, \tag{1}$$

where

$$\phi = e^{-\beta\epsilon_{\rm cost}}\frac{Z}{Z^{\rm closed}}\,. \tag{2}$$

For appropriate parameters, $P_{\rm open}$ presents a sharp transition from $P_{\rm open} \approx 0$ to $P_{\rm open} \approx 1$ as the volume is increased. We approximate this dependence with a Hill function

$$P_{\rm open} = \frac{(V/V^*)^h}{(V/V^*)^h+1}\,, \tag{3}$$

where $h$ is the Hill coefficient. Comparing Eq. (2) and Eq. (3), we obtain

$$h = \left.\frac{d\ln\phi}{d\ln V}\right|_{V^*}\,. \tag{4}$$

We shall derive an explicit expression for $\phi$ by evaluating the partition functions. We then will use the result to compute $h$ using Eq. (4).

#### 1.2 Computation of the partition function using the transfer matrix method

We here compute the partition functions $Z$ and $Z^{\rm closed}$ for different boundary conditions using the transfer matrix method. The transfer matrices for our model are

$$\hat{T} = \begin{pmatrix} 1 & x & x_{\rm adp} \\ 1 & xy & x_{\rm adp} \\ 1 & xy & x_{\rm adp} \end{pmatrix} \qquad \hat{T}^{\rm closed} = \begin{pmatrix} 1 & x^{\rm closed} & x^{\rm closed}_{\rm adp} \\ 1 & x^{\rm closed}y & x^{\rm closed}_{\rm adp} \\ 1 & x^{\rm closed}y & x^{\rm closed}_{\rm adp} \end{pmatrix}\,, \tag{5}$$

where

$$\begin{aligned} x &= \frac{a_{\rm atp}-c_{\rm atp}}{K_{\rm ori}}, \quad & x_{\rm adp} &= \frac{a_{\rm adp}-c_{\rm adp}}{K_{\rm ori}}, \\ x^{\rm closed} &= \frac{a_{\rm atp}-c_{\rm atp}}{K^{\rm closed}_{\rm ori}}, \quad & x^{\rm closed}_{\rm adp} &= \frac{a_{\rm adp}-c_{\rm adp}}{K^{\rm closed}_{\rm ori}} \\ y &= e^{\beta J}\,, \end{aligned} \tag{6}$$

where $K_{\rm ori}$ and $K^{\rm closed}_{\rm ori}$ are the dissociation constant at the weak origin sites in the open and closed DNA conformation, respectively. Since the two transfer matrices have the same structure, we focus for simplicity on the one for the open conformation.

The two non-vanishing eigenvalues of the transfer matrix are

$$\lambda_\pm = \frac{1+xy+rx \pm \sqrt{(1+xy+rx)^2-4x(y-1)}}{2}\,, \tag{7}$$

where $r = x_{\rm adp}/x$. Both eigenvalues are increasing functions of $x$, and we have $\lambda_- < 1 < \lambda_+$.

The transfer matrix is diagonalized by a transformation $\hat{D} = \hat{U}\hat{T}\hat{U}^{-1}$, with

$$\hat{D} = \begin{pmatrix} 0 & 0 & 0 \\ 0 & \lambda_- & 0 \\ 0 & 0 & \lambda_+ \end{pmatrix}\,, \tag{8}$$

and

$$\hat{U} = \begin{pmatrix} -x_{\rm adp} & 1-\lambda_+ & 1-\lambda_- \\ 0 & 1 & 1 \\ 1 & 1 & 1 \end{pmatrix} \qquad \hat{U}^{-1} = \begin{pmatrix} 0 & -1 & 1 \\ \frac{-1}{\lambda_+-\lambda_-} & \frac{1+x_{\rm adp}-\lambda_-}{\lambda_+-\lambda_-} & \frac{-x_{\rm adp}}{\lambda_+-\lambda_-} \\ \frac{1}{\lambda_+-\lambda_-} & \frac{\lambda_+-x_{\rm adp}-1}{\lambda_+-\lambda_-} & \frac{x_{\rm adp}}{\lambda_+-\lambda_-} \end{pmatrix}\,. \tag{9}$$

We now consider three specific scenario, corresponding to different boundary conditions.

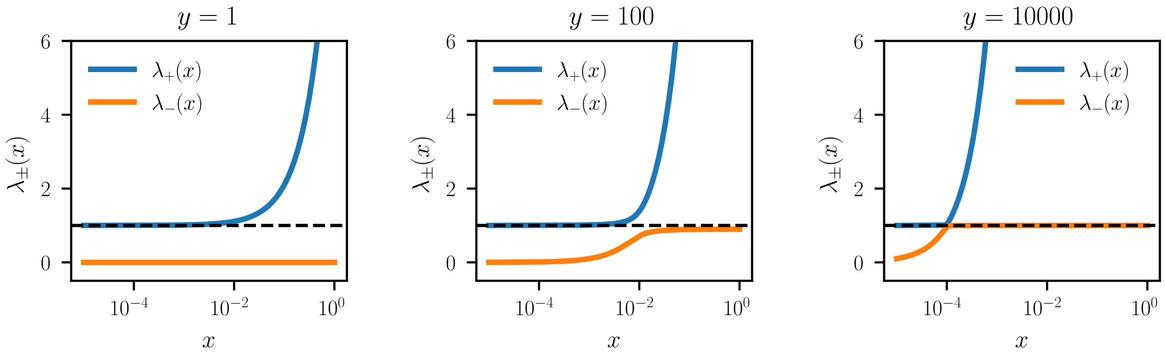

Figure 1: Eigenvalues of the transfer matrix as a function of $x$ for different values of the cooperativity weight $y$ assuming $r = 0$.

##### 1.3 Periodic cluster (ring) of weak sites

We consider a ring of $m$ weak sites. In this case, we have

$$Z = \mathrm{Tr}\,\hat{T}^m = (\lambda_+)^m + (\lambda_-)^m \sim (\lambda_+)^m \,. \tag{10}$$

The approximation comes from the observation that $\lambda_+$ is on the order of $x(y+r)$, while $\lambda_-$ is on the order of $y/[x(y+r)^2]$. This result implies that

$$\phi = e^{-\beta\epsilon_{\mathrm{cost}}} \left(\frac{\lambda_+}{\lambda_+^{\mathrm{closed}}}\right)^m \,. \tag{11}$$

We now discuss whether neglecting the contribution from the second eigenvalue is appropriate. In the case of strong cooperativity, the two eigenvalues are close to each other for a certain value of $x$, see Fig. 1). In contrast, in the absence of cooperativity ($y = 1$), $\lambda_- = 0$ and $\lambda_+ = 1 + x$.

##### 1.4 Linear cluster of weak sites

The origin is not a ring but a linear sequence of binding sites. For the sake of simplicity, we assume that the origin is formed by a single linear cluster of $m$ sites. Extension to a more detailed architecture that includes several smaller clusters relatively far apart would not significantly change the main results of this work. We consider a single linear cluster of $m$ sites. The partition function is given by

$$Z = \mathbf{v}^{\mathrm{T}} \hat{U} \hat{D}^m \hat{U}^{-1} \mathbf{u} \,. \tag{12}$$

with

$$\mathbf{v} = \begin{pmatrix} 1 \\ 0 \\ 0 \end{pmatrix} \qquad \mathbf{u} = \begin{pmatrix} 1 \\ 1 \\ 1 \end{pmatrix} \,. \tag{13}$$

The first matrix multiplications are

$$\hat{U}^{\mathrm{T}} \mathbf{v} = \begin{pmatrix} -x_{\mathrm{adp}} \\ 1 - \lambda_+ \\ 1 - \lambda_- \end{pmatrix} \qquad \hat{U}^{-1} \mathbf{u} = \begin{pmatrix} 0 \\ \frac{-\lambda_-}{\lambda_+ - \lambda_-} \\ \frac{\lambda_+}{\lambda_+ - \lambda_-} \end{pmatrix} \,. \tag{14}$$

We plug these expressions into Eq. (12), obtaining

$$Z = \frac{1}{\lambda_+ - \lambda_-} \left[ -\lambda_-^{m+1} \left[1 - \lambda_+\right] + \lambda_+^{m+1} \left[1 - \lambda_-\right] \right] \,. \tag{15}$$

##### 1.5 Linear cluster of weak sites flanked by a strong docking site

We consider a linear cluster having a strong site to its side. In this case, $m$ represents the number of weak sites in the cluster. The partition function is still given by Eq. (12), but the boundary vectors $\mathbf{v}$ and $\mathbf{u}$ need to be modified:

$$\mathbf{v} = \begin{pmatrix} 1 \\ x_{\text{dock}} \\ x_{\text{dock,adp}} \end{pmatrix} \qquad \mathbf{u} = \begin{pmatrix} 1 \\ 1 \\ 1 \end{pmatrix} . \tag{16}$$

Here, $x_{\text{dock}} = (a_{\text{atp}} - c_{\text{atp}})/K_{\text{dock}}$ and $x_{\text{dock,adp}} = (a_{\text{adp}} - c_{\text{adp}})/K_{\text{dock}}$, where $K_{\text{dock}}$ is the dissociation constant associated with the docking site. We assume that the dissociation constant for the docking site is the same in the two conformations, so that we can measure specifically how the docking site stabilizes the subsequent occupation of the adjacent weaker sites.

We now compute

$$\hat{U}^{\text{T}}\mathbf{v} = \begin{pmatrix} -x_{\text{adp}} + x_{\text{dock,adp}} \\ 1 - \lambda_+ + x_{\text{dock}} + x_{\text{dock,adp}} \\ 1 - \lambda_- + x_{\text{dock}} + x_{\text{dock,adp}} \end{pmatrix} \qquad \hat{U}^{-1}\mathbf{u} = \begin{pmatrix} 0 \\ \frac{-\lambda_-}{\lambda_+ - \lambda_-} \\ \frac{\lambda_+}{\lambda_+ - \lambda_-} \end{pmatrix} . \tag{17}$$

We plug these expressions into Eq. (12), obtaining

$$Z = \frac{1}{\lambda_+ - \lambda_-} \left[ -\lambda_-^{m+1} \left[1 - \lambda_+ + x_{\text{dock}} + x_{\text{dock,adp}}\right] + \lambda_+^{m+1} \left[1 - \lambda_- + x_{\text{dock}} + x_{\text{dock,adp}}\right] \right] . \tag{18}$$

When there is no cooperativity ($y = 1$), we have $\lambda_- = 0$ and the correction of the docking sites provides a similar prefactor in the term for the open and the closed conformation:

$$Z \simeq \lambda_+^m (1 + x_{\text{dock}} + x_{\text{dock,adp}}) \simeq \lambda_+^m (x_{\text{dock}} + x_{\text{dock,adp}}) . \tag{19}$$

In the presence of cooperativity, all terms in the right hand side of Eq. (18) contribute in the equation for the open conformation.

Conversely, for the closed conformation we can still use the approximation $x^{\text{closed}} \ll 1$ and thus

$$Z^{\text{closed}} \simeq (\lambda_+^{\text{closed}})^m (x_{\text{dock}}^{\text{closed}} + x_{\text{dock,adp}}^{\text{closed}}) . \tag{20}$$

### 2 Calculation of the Hill coefficient

#### 2.1 Calculation of $x^*$

We calculate the midpoint volume $V^*$ and then use Eq. (4) to find the Hill coefficient $h$. We start by finding the midpoint value of $x$, $x = x^*$, see Eq. (6).

We define

$$\eta = \frac{x}{x^{\text{closed}}} = \frac{K_{\text{ori}}^{\text{closed}}}{K_{\text{ori}}} , \tag{21}$$

so that Eq. (2) becomes

$$\phi^{1/m} = e^{-\frac{\beta\epsilon_{\text{cost}}}{m}} \frac{1 + xy + \sqrt{(1 + xy + x_{\text{adp}})^2 - 4x(y-1)}}{1 + \frac{x}{\eta}y + \sqrt{(1 + \frac{x}{\eta}y + \frac{x_{\text{adp}}}{\eta})^2 - 4\frac{x}{\eta}(y-1)}} . \tag{22}$$

We assume that $\eta$ is large, so that the expression reduces to

$$\begin{aligned} \phi^{1/m} &= \frac{e^{-\beta\epsilon_{\text{cost}}/m}}{2} \left[ 1 + yx + x_{\text{adp}} + \sqrt{(1 + yx + x_{\text{adp}})^2 - 4x(y-1)} \right] \\ &= \frac{e^{-\beta\epsilon_{\text{cost}}/m}}{2} \left[ 1 + (y + r)x + \sqrt{[1 + (y+r)x]^2 - 4x(y-1)} \right] . \end{aligned} \tag{23}$$

We expect to have more free DnaA-ATP than DnaA-ADP at initiation, so that $r$ should be less than 1 for $V = V^*$. We also define $A = e^{-\beta\epsilon_{\text{cost}}/m}$ for convenience of notation.

We now impose $\phi = 1$ for $x = x^*$, obtaining

$$\frac{2}{A} - [1 + (y + r)x^*] = \sqrt{[1 + (y + r)x^*]^2 - 4x^*(y-1)} . \tag{24}$$

Squaring both sides of the equation we get

$$\frac{4}{A^2} - \frac{4}{A}[1 + (y + r)x^*] = -4x^*(y - 1)\,, \tag{25}$$

which simplifies to

$$1 - A[1 + (y + r)x^*] = -A^2 x^*(y - 1)\,, \tag{26}$$

so that

$$x^* = \frac{1 - A + A^2}{A(y + r) - A^2 y}. \tag{27}$$

We now assume that $y \gg r$ and, since $A < 1$, we neglect the $A^2$ in the numerator, obtaining

$$x^* = \frac{1}{Ay} = e^{\beta\left(\frac{\epsilon_{\text{cost}}}{m} - J\right)}. \tag{28}$$

This calculation shows that, under our assumptions, the value $x = x^*$ is unaffected by the presence of DnaA-ADP, because the concentration of DnaA-ADP is not large enough to initiate replication without cooperative binding.

#### 2.2 Equilibrium occupancy of the chromosomal sites

The concentrations of occupied sites are given by detailed balance:

$$\begin{aligned} K\,c_{\text{atp}} &= (a_{\text{atp}} - c_{\text{atp}})(c_{\text{tot}} - c_{\text{atp}} - c_{\text{adp}})\,, \\ K\,c_{\text{adp}} &= (a_{\text{adp}} - c_{\text{adp}})(c_{\text{tot}} - c_{\text{atp}} - c_{\text{adp}})\,. \end{aligned} \tag{29}$$

Summing the two equations in Eq. (29), we obtain

$$Kc = (a - c)(c_{\text{tot}} - c)\,, \tag{30}$$

where $c = c_{\text{atp}} + c_{\text{adp}}$ . The solution of this second order algebraic equation is

$$c = \frac{a + c_{\text{tot}} + K - \sqrt{(a + c_{\text{tot}} + K)^2 - 4ac_{\text{tot}}}}{2}\,. \tag{31}$$

The proportion of sites bound to DnaA-ATP and DnaA-ADP are

$$c_{\text{atp}} = \alpha c\,, \qquad c_{\text{adp}} = (1 - \alpha)c\,, \tag{32}$$

where $\alpha = a_{\text{atp}}/a$. Indeed, looking for solutions of the form $c_{\text{atp}} = \xi c$ in the first equation, we obtain

$$K\xi c = (\alpha a - \xi c)(c_{\text{tot}} - c)\,, \tag{33}$$

and we need $\alpha = \xi$ for Eq. (33) to be compatible with Eq. (30). Equation (32) is a consequence of the fact that the dissociation constant for the sites is assumed to be the same for the two nucleotide states of DnaA, therefore the partitioning of the proteins bound to the chromosomal sites matches the total proportions of proteins in the cell.

#### 2.3 Volume at midpoint at fixed $\alpha$

We now find an explicit expression for $V^*$ at fixed $\alpha$. We define

$$z = K_{\text{ori}} x^* \simeq K_{\text{ori}} e^{\beta\left(\frac{\epsilon_{\text{cost}}}{m} - J\right)}\,. \tag{34}$$

Given the definition of $x$, $z$ is equal to the concentration of free DnaA-ATP at initiation:

$$z = (a_{\text{atp}} - c^*_{\text{atp}}) = \alpha(a - c^*)\,. \tag{35}$$

Combining Eq. (34) with Eq. (35) we obtain the condition for the midpoint

$$K_{\text{ori}} e^{\beta\left(\frac{\epsilon_{\text{cost}}}{m} - J\right)} = \alpha(a - c^*)\,, \tag{36}$$

where the left hand side depends on the origin parameters, while the right hand side contains the effect of titration.

Solving Eq. (31) in $c_{\text{tot}} = c^*_{\text{tot}}$ and using Eq. (35) to eliminate $c^*$ we obtain

$$c^*_{\text{tot}} = \frac{\alpha K a}{z} - K + a - \frac{z}{\alpha} = \frac{(\alpha a - z)(\alpha K + z)}{\alpha z}\,. \tag{37}$$

Therefore, the volume at the midpoint is expressed by

$$V^* = \frac{\alpha n^* z}{(\alpha a - z)(\alpha K + z)}\,, \tag{38}$$

where we have defined the number of titration sites at initiation $n^* = c^*_{\text{tot}} V^*$.

#### 2.4 Calculation of the Hill coefficient at fixed $\alpha$ and $n$

We calculate the Hill coefficient in case $h_{\mathrm{rep}} = h_{\mathrm{hyd}} = 0$, using Eq. (4) and the chain rule:

$$h = h_{\mathrm{dil}} = \left.\frac{d\ln\phi}{dx}\right|_{V^*} \left.\frac{dx}{dc}\right|_{V^*} \left.\frac{dc}{dc_{\mathrm{tot}}}\right|_{V^*} \left.\frac{dc_{\mathrm{tot}}}{d\ln V}\right|_{V^*} . \tag{39}$$

We calculate the derivatives appearing in the right hand side of Eq. (39) one by one. At the moment we keep the active fraction $\alpha$ and the total number of titration sites $n$ fixed. The first term in Eq. (39) is given by

$$\begin{aligned} \left.\frac{d\ln\phi}{dx}\right|_{x^*} &= \frac{me^{-\beta\epsilon_{\mathrm{cost}}/m}}{2} \left.\frac{d\lambda_+}{dx}\right|_{x^*} \\ &= \frac{me^{-\beta\epsilon_{\mathrm{cost}}/m}}{2} \left[(y+r) + \frac{(y+r)[1+(y+r)x^*] - 2(y-1)}{\sqrt{[1+(y+r)x^*]^2 - 4x^*(y-1)}}\right] \\ &= me^{-\beta\epsilon_{\mathrm{cost}}/m} \frac{e^{\beta\epsilon_{\mathrm{cost}}/m}(y+r) - (y-1)}{2e^{\beta\epsilon_{\mathrm{cost}}/m} - 1 - (y+r)x^*} \\ &\simeq me^{-\beta\epsilon_{\mathrm{cost}}/m} \frac{e^{\beta\epsilon_{\mathrm{cost}}/m}(y+r) - (y-1)}{e^{\beta\epsilon_{\mathrm{cost}}/m} - 1} \\ &\simeq mye^{-\beta\epsilon_{\mathrm{cost}}/m} . \end{aligned} \tag{40}$$

In the third line, we have used that $\phi^{1/m} = e^{-\beta\epsilon_{\mathrm{cost}}/m}\lambda_+ = 1$ at the midpoint, so that

$$\sqrt{[1+(y+r)x^*]^2 - 4x^*(y-1)} = 2e^{\beta\epsilon_{\mathrm{cost}}/m} - 1 - (y+r)x^* . \tag{41}$$

We now move to the second term. From the definition of $x$ in Eq. (6) and Eq. (32) we immediately obtain

$$x = \alpha\frac{(a-c)}{K_{\mathrm{ori}}} . \tag{42}$$

We therefore have

$$\frac{dx}{dc} = -\frac{\alpha}{K_{\mathrm{ori}}} . \tag{43}$$

For the third term, using Eq. (31) we have

$$\left.\frac{dc}{dc_{\mathrm{tot}}}\right|_{c^*_{\mathrm{tot}}} = \frac{1}{2}\left[1 - \frac{(c^*_{\mathrm{tot}} + K - a)}{\sqrt{(a + c^*_{\mathrm{tot}} + K)^2 - 4ac^*_{\mathrm{tot}}}}\right] . \tag{44}$$

To simplify the square root term, we use again Eq. (31):

$$\begin{aligned} \sqrt{(a + c^*_{\mathrm{tot}} + K)^2 - 4ac^*_{\mathrm{tot}}} &= a + c^*_{\mathrm{tot}} + K - 2c^* \\ &= \frac{z}{\alpha} + c^*_{\mathrm{tot}} - c^* + K \\ &= \frac{z}{\alpha} + \frac{Kc^*}{a - c^*} + K \\ &= \frac{z}{\alpha} + \frac{Ka\alpha}{z} , \end{aligned} \tag{45}$$

where we have used Eqs. (30) and (35). Substituting this result into Eq. (44) we obtain

$$\begin{aligned} \left.\frac{dc}{dc_{\mathrm{tot}}}\right|_{c^*_{\mathrm{tot}}} &= \frac{1}{2}\left[1 - \frac{(c^*_{\mathrm{tot}} + K - a)}{\frac{z}{\alpha} + \frac{Ka\alpha}{z}}\right] \\ &= \frac{1}{2}\left[1 - \frac{\frac{Ka\alpha}{z} - \frac{z}{\alpha}}{\frac{z}{\alpha} + \frac{Ka\alpha}{z}}\right] \\ &= \frac{z/\alpha}{z/\alpha + Ka\alpha/z} . \end{aligned} \tag{46}$$

Finally, since $c_{\mathrm{tot}} = n/V$, we have

$$\left.\frac{dc_{\mathrm{tot}}}{d\ln V}\right|_{V^*} = -c^*_{\mathrm{tot}} = -\frac{(\alpha a - z)(\alpha K + z)}{\alpha z} . \tag{47}$$

Substituting all of these results into Eq. (39), we have

$$
\begin{aligned}
h &= mye^{-\beta\epsilon_{\mathrm{cost}}/m}\frac{\alpha}{K_{\mathrm{ori}}}\frac{z/\alpha}{z/\alpha+K a\alpha/z}\frac{(\alpha a-z)(\alpha K+z)}{\alpha z}\\
&= m\frac{1}{z/\alpha+K a\alpha/z}\frac{(\alpha a-z)(\alpha K+z)}{\alpha z}\\
&= m\left(a-\frac{z}{\alpha}\right)\left(\frac{K\alpha}{z}+1\right)\left(\frac{1}{z/\alpha+K a\alpha/z}\right), \qquad (48)
\end{aligned}
$$

and, defining the ratio $w = z/\alpha$:

$$
h(w) = m\,(a-w)\left(\frac{K}{w}+1\right)\left(\frac{1}{w+Ka/w}\right). \qquad (49)
$$

#### 2.5 Optimal Hill coefficient at fixed $\alpha$ and $n$

We look for the value of $w$ that maximizes the Hill coefficient:

$$
\frac{dh}{dw} = m\left[\frac{a-w}{w^2+Ka}-\frac{w+K}{w^2+Ka}-\frac{2w(w+K)(a-w)}{(w^2+Ka)^2}\right]=0\,, \qquad (50)
$$

obtaining

$$
\begin{aligned}
0 &= (a-w)(w^2+Ka)-(w+K)(w^2+Ka)-2w(w+K)(a-w) && (51)\\
&= aw^2+Ka^2-w^3-Kaw-w^3-Kaw-Kw^2-K^2a-2aw^2+2w^3-2Kaw+2Kw^2 && (52)\\
&= -w^2(a-K)-4Kaw+Ka^2-K^2a\,. && (53)
\end{aligned}
$$

Using that $a \gg K$, we obtain

$$
w_{\mathrm{opt}} = \frac{2Ka-\sqrt{4(Ka)^2+Ka^3}}{-a} \simeq \sqrt{Ka}\,. \qquad (54)
$$

This is also the value of $w$ for which the total concentration of initiator proteins is equal to the concentration of binding sites. Indeed, if we consider Eq. (37) with the condition

$$
c^*_{\mathrm{tot}} = a\,, \qquad (55)
$$

we obtain

$$
\frac{Ka}{w}-w-K=0\,, \qquad (56)
$$

and thus

$$
w = \frac{-K+\sqrt{K^2+4Ka}}{2} \simeq \sqrt{Ka}\,. \qquad (57)
$$

Substituting this value back into Eq. (49) we estimate the optimal value of the Hill coefficient

$$
h_{\mathrm{opt}} = m\left(1+\sqrt{\frac{K}{a}}\right)\left(\frac{a-\sqrt{Ka}}{2\sqrt{Ka}}\right) \simeq \frac{m}{2}\sqrt{\frac{a}{K}}\,. \qquad (58)
$$

#### 2.6 Hill coefficient for varying $\alpha$ and $n$

In general, the Hill coefficient can be decomposed as

$$
h = \overbrace{\frac{\partial\ln\phi}{\partial c_{\mathrm{tot}}}\frac{\partial c_{\mathrm{tot}}}{\partial\ln V}\bigg|_{V^*}}^{h_{\mathrm{dil}}} + \overbrace{\frac{\partial\ln\phi}{\partial c_{\mathrm{tot}}}\frac{\partial c_{\mathrm{tot}}}{\partial n}\frac{dn}{d\ln V}\bigg|_{V^*}}^{h_{\mathrm{rep}}} + \overbrace{\frac{\partial\ln\phi}{\partial\alpha}\frac{\partial\alpha}{\partial\ln V}\bigg|_{V^*}}^{h_{\mathrm{hyd}}}. \qquad (59)
$$

We have already derived $h_{\mathrm{dil}}$ in the subsection 2.4. We now derive the other two terms.

#### 2.7 Calculation of $h_{\rm hyd}$

In the model, the fraction $\alpha$ is determined by

$$\alpha = \max\left[\alpha_{\rm min}\,,\,\alpha_{\rm max}\frac{V-\chi_0 n_{\rm forks}}{V}\right]\,, \tag{60}$$

where $n_{\rm forks}$ is the number of replication forks, and $\alpha_{\rm min}$, $\alpha_{\rm max}$ and $\chi_0$ are positive constants. We have

$$\frac{\partial\alpha}{\partial\ln V} = \alpha_{\rm max}-\alpha \quad\longrightarrow\quad 0\leq\frac{\partial\alpha}{\partial\ln V}\leq\alpha_{\rm max}\,. \tag{61}$$

In slow growth, before initiation one has $n_{\rm forks}=0$, so that $\alpha=\alpha_{\rm max}$ and $\partial\alpha/\partial\ln V=0$.

We now return to the general case. Since $r=(1-\alpha)/\alpha$, we have

$$\left.\frac{\partial\ln\phi}{\partial\alpha}\right|_{x^*} = -\frac{1}{\alpha^2}\left.\frac{\partial\ln\phi}{\partial r}\right|_{x^*} + \left.\frac{\partial\ln\phi}{\partial x}\frac{\partial x}{\partial\alpha}\right|_{x^*}\,. \tag{62}$$

We start by the first term:

$$\begin{aligned}
-\frac{1}{\alpha^2}\left.\frac{\partial\ln\phi}{\partial r}\right|_{x^*} &= -\frac{1}{\alpha^2}\frac{m}{\lambda_+(x^*)}\left.\frac{d\lambda_+}{dr}\right|_{x^*}\\
&= -\frac{me^{-\beta\epsilon_{\rm cost}/m}}{2\alpha^2}\left[\frac{x^*}{2}+\frac{2x^*[1+x^*(y+r)]}{4\sqrt{[1+(y+r)x^*]^2-4x^*(y-1)}}\right]\\
&= -\frac{me^{-\beta\epsilon_{\rm cost}/m}}{2\alpha^2}\frac{x^*}{2}\left[1+\frac{1+x^*(y+r)}{\sqrt{[1+(y+r)x^*]^2-4x^*(y-1)}}\right]\\
&= -\frac{me^{-\beta\epsilon_{\rm cost}/m}}{2\alpha^2}\frac{x^*}{2}\left[1+\frac{1+x^*(y+r)}{2e^{\beta\epsilon_{\rm cost}/m}-1-x^*(y+r)}\right]\\
&= -\frac{me^{-\beta\epsilon_{\rm cost}/m}}{2\alpha^2}\frac{x^*}{2}\left[\frac{2e^{\beta\epsilon_{\rm cost}/m}}{2e^{\beta\epsilon_{\rm cost}/m}-1-x^*(y+r)}\right]\\
&= -\frac{me^{-\beta\epsilon_{\rm cost}/m}}{2\alpha^2 y}\left[\frac{1}{2e^{\beta\epsilon_{\rm cost}/m}-1-\frac{e^{-\beta\epsilon_{\rm cost}/m}}{y}(y+r)}\right]\,.
\end{aligned} \tag{63}$$

We now move to the second term:

$$\begin{aligned}
\left.\frac{\partial\ln\phi}{\partial x}\frac{\partial x}{\partial\alpha}\right|_{x^*} &= mye^{-\beta\epsilon_{\rm cost}/m}\frac{x^*}{\alpha}\\
&= \frac{m}{\alpha}\,.
\end{aligned} \tag{64}$$

We now use these results to empirically estimate $h_{\rm hyd}$ for *E. coli* in optimal growth. Realistic parameters are $y\approx 50$, $\beta\epsilon_{\rm cost}\in(10,20)$, $m\approx 8$, $\alpha\approx 0.15$, which give

$$-\frac{1}{\alpha^2}\left.\frac{\partial\ln\phi}{\partial r}\right|_{x^*} < 10\,, \tag{65}$$

and

$$\left.\frac{\partial\ln\phi}{\partial x}\frac{\partial x}{\partial\alpha}\right|_{x^*} \approx 53 \tag{66}$$

so that the second term dominates over the first. We also have $\partial\alpha/\partial\ln V\approx 0.85$, so that

$$h_{\rm hyd}\approx 45\,. \tag{67}$$

#### 2.8 Calculation of $h_{\rm rep}$

The number of chromosomal sites varies as

$$\frac{dn}{dt} = n_{\rm forks}v\rho\,, \tag{68}$$

where $n_{\text{forks}}$ is the number of active forks, $v$ is their speed, and $\rho$ is the density of chromosomal sites along the genome. Given that $c_{\text{tot}} = n/V$, we have

$$\frac{\partial c_{\text{tot}}}{\partial n} \frac{dn}{d\ln V}\bigg|_{V^*} = \frac{1}{V^*} \frac{dn}{d\ln V}\bigg|_{V^*} . \tag{69}$$

Using the dependence of the volume on the growth rate, $\Lambda dt = d\ln V$, we simply have

$$\frac{dn}{d\ln V} = \frac{n_{\text{forks}} v\rho}{\Lambda} . \tag{70}$$

Using the fact that

$$\frac{d\ln\phi}{dc_{\text{tot}}} = \frac{h_{\text{dil}}}{\frac{dc_{\text{tot}}}{d\ln V}} , \tag{71}$$

we have

$$\frac{\partial \ln\phi}{\partial c_{\text{tot}}}\bigg|_{V^*} = -\frac{h_{\text{dil}}}{c^*_{\text{tot}}} , \tag{72}$$

and thus

$$h_{\text{rep}} = -\frac{h_{\text{dil}}}{c^*_{\text{tot}}} \frac{n_{\text{forks}} v\rho}{\Lambda V^*} = -h_{\text{dil}} \frac{n_{\text{forks}} v\rho}{\Lambda n^*} . \tag{73}$$

We estimate $h_{\text{rep}}$ for E.coli in optimal growth conditions ($\tau = 25$ min). We first determine whether initiation takes place before or after termination. The time of initiation in the regime $\tau < C < 2\tau$ is given by

$$t_{\text{init}} = \begin{cases} 2\tau - D - C \equiv t_{\text{ter}} - (C - \tau) & \text{if } 2\tau > D + C , \\ 3\tau - D - C \equiv t_{\text{ter}} + (2\tau - C) & \text{if } 2\tau < D + C . \end{cases} \tag{74}$$

We assume $D = 20$ min, so that $2\tau = 50 < D + C$. Therefore, the time of termination is $t_{\text{ter}} = \tau - D = 5$ min and the time of initiation is $t_{\text{init}} = 3\tau - D - C = 15$ min. There are two genomes in the cell at the time of initiation. The replication progress between two initiations is $\tau/C$ in steady state. Therefore, if the total number of sites per chromosome is $n_{\text{sites}} = 300$, then

$$n^* = 2n_{\text{sites}} \left[1 + \frac{\tau}{C}\right] = 600 \left(1 + \frac{25}{40}\right) = 975 . \tag{75}$$

Before initiation, we have

$$\frac{n_{\text{forks}} v\rho}{\Lambda} = \frac{4 \cdot 1000 \,\text{bp/s} \ \cdot 300 \cdot (25 \cdot 60 \,\text{s})}{4 \cdot 10^6 \,\text{bp} \cdot \ln 2} = 649 , \tag{76}$$

and thus

$$n^* > n_{\text{forks}} v\rho , \tag{77}$$

right before initiation. Then, after the initiation, the number of forks moves from $n_{\text{forks}} = 4$ to $n_{\text{forks}} = 12$ and

$$n_{\text{forks}} v\rho \to 1947 , \tag{78}$$

and thus

$$n^* < n_{\text{forks}} v\rho , \tag{79}$$

right after initiation.

Now we make an estimate of $h_{\text{rep}}$ in these conditions, under the assumptions made for Eq.(67). Before initiation, we have

$$h_{\text{rep}} = -\frac{649}{975} h_{\text{dil}} \approx -0.66\, h_{\text{dil}} . \tag{80}$$

Therefore, if $h_{\text{dil}} = 100$, the value of the total Hill coefficient in fast growth becomes

$$h \approx 100 - 66 + 45 = 79 , \tag{81}$$

where we have used Eq. (67) for the contribution $h_{\text{hyd}}$.

#### 3 Stability analysis

##### 3.1 Midpoint volume at varying $\alpha$

We derive a formula for $V^*$ that, in contrast with Eq. (38), explicitly takes into account that $\alpha$ depends itself on $V$, see Eq. (60). Close to initiation, when $\alpha$ is growing, we have

$$\alpha(V) = \alpha_{\max}\left[1 - \frac{n_{\text{forks}}\chi_0}{V}\right] \equiv \alpha_{\max}\left[1 - \frac{\chi}{V}\right], \tag{82}$$

where we defined $\chi = n_{\text{forks}}\chi_0$. The expression for the midpoint volume becomes

$$V^* = \frac{\alpha(V^*)n^*z}{(\alpha(V^*)a - z)(\alpha(V^*)K + z)} \simeq \frac{\alpha(V^*)n^*}{\alpha(V^*)a - z}, \tag{83}$$

where the approximation follows from the fact that $K$ is small. Solving for $V^*$, and keeping only the biologically admissible solution for which $\alpha > 0$, we obtain

$$V^* = \frac{n^* + a\chi + \sqrt{[n^* - a\chi]^2 + \frac{4n^*\chi z}{\alpha_{\max}}}}{2\left(a - \frac{z}{\alpha_{\max}}\right)}. \tag{84}$$

We now use the facts that $V^*$ is close to $n^*/a$ at initiation and that $\alpha$ is a small number. Therefore, we expand $n^*$ around $n^* = a\chi$ to obtain

$$V^* = \frac{\frac{n^*}{a} + \chi}{2\left[1 - \sqrt{\frac{z}{a\alpha_{\max}}}\right]}. \tag{85}$$

We note that the affinity for the origin binding sites should always be such that $z < a\alpha_{\max}$. Since $z$ is the concentration of free DnaA-ATP at the midpoint, this constraint means that the maximal concentration of DnaA-ATP must be large enough to trigger initiation.

##### 3.2 Conditions of stability in case of multi-fork replication

A sharp response is not enough to ensure proper timing of replication. In case of multi-fork replication, small deviations from the proper time interval between initiation events might be amplified from generation to generation even when the firing rate is a step function of the volume. In this section we perform a stability analysis assuming stepwise response at $V^*$, to determine the conditions on the RIDA mechanism to ensure precise timing of initiation. A similar analysis in the absence of hydrolysis was first proposed by Fu et al. [1] to show that the system is not stable in the regime $\tau < C < 2\tau$, where $\tau$ is the doubling time and $C$ the replication time.

Initiation is triggered whenever

$$V = V^*. \tag{86}$$

We need to find the relation between two subsequent intervals between consecutive initiations. Let $t_a$ be the time between two initiations and $t_b$ the time to be waited for a third one afterward. By construction, we have

$$V(t_a) = V^*(t_a), \tag{87}$$

and

$$V(t_b) = V^*(t_b). \tag{88}$$

Moreover, we also use the fact that $V$ increases in time exponentially at rate $\Lambda$, whereas $V^*$ evolves according to Eq. (85). In the regime $\tau < C < 2\tau$, for every genome the number of sites right before the first initiation is given by the sum of the sites of one full chromosome and the newly created sites during the time $t_a$:

$$n = n_{\text{gen}}n_{\text{sites}}\left[1 + \frac{t_a}{C}\right], \tag{89}$$

where $n_{\text{sites}}$ is the number of sites of one full chromosome and $n_{\text{gen}}$ is the number of genomes in the cell right before initiation. We have

$$n_{\text{gen}} = \begin{cases} 1 & \text{if } t_{\text{init}} < t_{\text{ter}} \\ 2 & \text{if } t_{\text{init}} > t_{\text{ter}}. \end{cases} \tag{90}$$

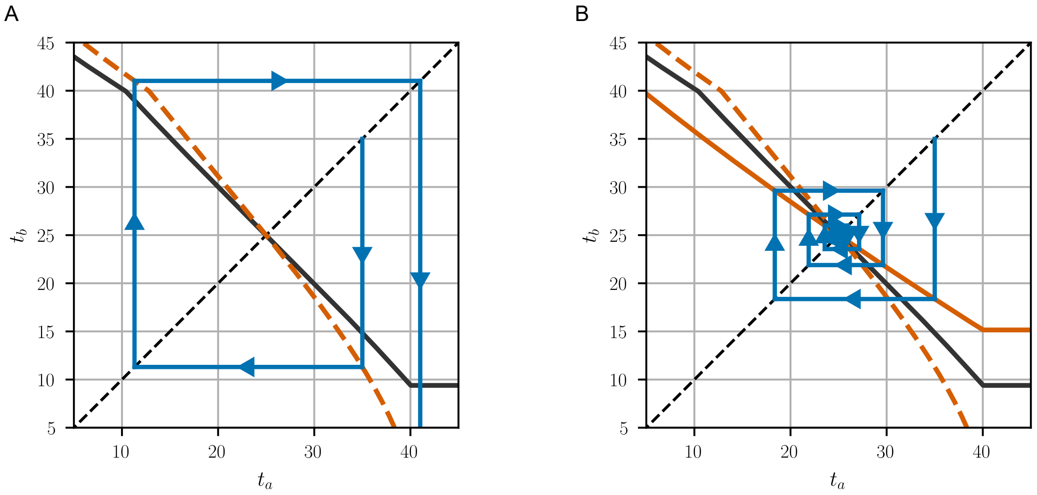

Figure 2: Cobweb plot for consecutive intervals between initiation events in case $\gamma := a\frac{2\chi_0}{n_{\rm sites}} = 0$ (A) and $\gamma = 0.6$ (B). The orange lines are given by Eq.(92) in case $\gamma = 0$ (dashed) and $\gamma = 0.6$ (continuous). The black continuous line was obtained using the threshold value for $\chi_0$, expressed in Eq.(94).

The total number of forks in the regime $\tau < C < 2\tau$ is

$$n_{\rm forks} = 2n_{\rm gen}\,. \tag{91}$$

We can now use Eq. (87) and Eq. (88) to find the relation between two consecutive intervals between initiation events:

$$e^{\Lambda t_b}\left[\left[1+\frac{t_a}{C}\right]+a\frac{2\chi_0}{n_{\rm sites}}\right] = 2\left[\left[1+\frac{t_b}{C}\right]+a\frac{2\chi_0}{n_{\rm sites}}\right]\,. \tag{92}$$

The stability condition is

$$\left|\frac{dt_b}{dt_a}(t_a = t_b = \tau)\right| < 1\,, \tag{93}$$

resulting in

$$\chi_0 > \frac{n_{\rm sites}}{a}\left[\left[\frac{2}{\ln 2}-1\right]\frac{\tau}{C}-1\right]\,. \tag{94}$$

Only when Eq. (94) is satisfied the intervals between initiations converge (Fig.2).

### 4 Special chromosomal sites

In this section, we consider a variant of the model in which DnaA-ATP has access to sites that are not accessible to DnaA-ADP, but with the same dissociation constant $K$. We use the index 1 to indicate the ordinary sites that can be occupied by DnaA in either state and the index 2 for the other special sites. We call $c_{\rm tot}$ the total concentrations of the ordinary sites and

$$c_{2,\rm tot} = \theta c_{\rm tot}\,, \tag{95}$$

the total concentration of special sites, where $\theta$ is a constant.

The chemical balance reads

$$\begin{aligned} K\,c_{1,\rm atp} &= (a_{\rm atp} - c_{1,\rm atp} - c_{2,\rm atp})(c_{\rm tot} - c_{1,\rm atp} - c_{1,\rm adp})\,,\\ K\,c_{2,\rm atp} &= (a_{\rm atp} - c_{1,\rm atp} - c_{2,\rm atp})(\theta c_{\rm tot} - c_{2,\rm atp})\,,\\ K\,c_{1,\rm adp} &= (a_{\rm adp} - c_{1,\rm adp})(c_{\rm tot} - c_{1,\rm atp} - c_{1,\rm adp})\,. \end{aligned} \tag{96}$$

where the index for the concentration of occupied binding sites represents the nucleotide state of the protein bound to the given binding site, therefore, for example, $c_{1,\rm adp}$ is the concentration of ordinary sites bound by DnaA-ADP.

We assume that the special sites are few compared to the ordinary sites, so that we can look for solutions for small $\theta$:

$$c_{1,\text{atp}} = c^{(0)}_{1,\text{atp}} + \theta c^{(1)}_{1,\text{atp}}\,, \qquad c_{1,\text{adp}} = c^{(0)}_{1,\text{adp}} + \theta c^{(1)}_{1,\text{adp}}\,, \qquad c_{2,\text{atp}} = \theta c^{(1)}_{2,\text{atp}}\,. \tag{97}$$

The solutions are found by plugging these equations into Eqs. (96) and by matching orders in $\theta$. For the zeroth order $c^{(0)}_1 := c^{(0)}_{1,\text{atp}} + c^{(0)}_{1,\text{adp}}$ we use the result we obtained in the previous sections:

$$c^{(0)}_1 = \frac{a + c_{\text{tot}} + K - \sqrt{(a + c_{\text{tot}} + K)^2 - 4ac_{\text{tot}}}}{2}\,. \tag{98}$$

Then, we simply have $c^{(0)}_{1,\text{atp}} = \alpha c^{(0)}_1$ and $c^{(0)}_{1,\text{adp}} = (1-\alpha)c^{(0)}_1$.

We now proceed with the first order terms. We start with the occupation of special sites $c_{2,\text{atp}}$. The equation for the first order in $\theta$ is

$$K\, c^{(1)}_{2,\text{atp}} = (a_{\text{atp}} - c^{(0)}_{1,\text{atp}})(c_{\text{tot}} - c^{(1)}_{2,\text{atp}})\,, \tag{99}$$

and the solution is

$$c_{2,\text{atp}} = \theta c^{(1)}_{2,\text{atp}} = \theta c_{\text{tot}} \left[1 - \frac{K}{a_{\text{atp}} - c^{(0)}_{1,\text{atp}} + K}\right] \equiv \theta c_{\text{tot}} \left[1 - \frac{K}{\alpha(a - c^{(0)}_1) + K}\right]\,. \tag{100}$$

As expected, the concentration of occupied special sites is closer to their total number the smaller is $K$ compared to the concentration of free proteins. Before we find $c^{(1)}_{1,\text{atp}}$ and $c^{(1)}_{1,\text{adp}}$, it is useful to obtain their sum:

$$c^{(1)}_1 := c^{(1)}_{1,\text{atp}} + c^{(1)}_{1,\text{adp}} = -\frac{c^{(1)}_{2,\text{atp}}\left[c_{\text{tot}} - c^{(0)}_1\right]}{K + (a - c^{(0)}_1) + (c_{\text{tot}} - c^{(0)}_1)} \equiv -\chi c^{(1)}_{2,\text{atp}}\,, \tag{101}$$

where we have defined $\chi$ to separate $c^{(1)}_{2,\text{atp}}$ from the rest of the expression. Note that $c^{(1)}_1$ is always negative, because the addition of the special sites reduces the available pool of proteins. The equations for the remaining concentrations $c^{(1)}_{1,\text{atp}}$ and $c^{(1)}_{1,\text{adp}}$ can be solved in terms of $c^{(1)}_1$. In the case of DnaA-ADP we have

$$c^{(1)}_{1,\text{adp}} = -\frac{(a_{\text{adp}} - c^{(0)}_{1,\text{adp}})c^{(1)}_1}{K + c_{\text{tot}} - c^{(0)}_1} = \frac{(a_{\text{adp}} - c^{(0)}_{1,\text{adp}})\left|c^{(1)}_1\right|}{K + c_{\text{tot}} - c^{(0)}_1}\,, \tag{102}$$

which is always positive because, thanks to the special sites titrating more DnaA-ATP, the DnaA-ADP proteins have less competition for the ordinary sites, and

$$\begin{aligned} c^{(1)}_{1,\text{atp}} &= -\frac{c^{(1)}_{2,\text{atp}}\left[c_{\text{tot}} - c^{(0)}_1\right]}{K + c_{\text{tot}} - c^{(0)}_1}\left[1 - \frac{a_{\text{atp}} - c^{(0)}_{1,\text{atp}}}{K + (a - c^{(0)}_1) + (c_{\text{tot}} - c^{(0)}_1)}\right] && (103)\\ &= c^{(1)}_1\left[1 + \frac{(a_{\text{adp}} - c^{(0)}_{1,\text{adp}})}{K + c_{\text{tot}} - c^{(0)}_1}\right] && (104)\\ &= c^{(1)}_1\left[1 + (1-\alpha)\frac{(a - c^{(0)}_1)}{K + c_{\text{tot}} - c^{(0)}_1}\right]\,. && (105)\end{aligned}$$

The total concentration of sites occupied by DnaA-ATP is

$$\begin{aligned} c_{\text{atp}} &= \alpha c^{(0)}_1 + \theta\left[c^{(1)}_{1,\text{atp}} + c^{(1)}_{2,\text{atp}}\right] && (106)\\ &= \alpha c^{(0)}_1 + \theta c^{(1)}_{2,\text{atp}}\left[1 - \chi\left[1 + (1-\alpha)\frac{a - c^{(0)}_1}{K + c_{\text{tot}} - c^{(0)}_1}\right]\right]\,. && (107)\end{aligned}$$

Writing explicitly $\chi$ and $c^{(1)}_{2,\text{atp}}$, we obtain

$$\begin{aligned} c_{\text{atp}} =& \alpha c^{(0)}_1 + \theta c_{\text{tot}}\left[1 - \frac{K}{\alpha(a - c^{(0)}_1) + K}\right] \times \\ &\times \left[1 - \frac{c_{\text{tot}} - c^{(0)}_1}{K + (a - c^{(0)}_1) + (c_{\text{tot}} - c^{(0)}_1)}\left[1 + (1-\alpha)\frac{a - c^{(0)}_1}{K + c_{\text{tot}} - c^{(0)}_1}\right]\right] && (108)\\ &\equiv \alpha c^{(0)}_1 + c_{\text{tot}}\omega\left[1 - \frac{c_{\text{tot}} - c^{(0)}_1}{K + (a - c^{(0)}_1) + (c_{\text{tot}} - c^{(0)}_1)}\left[1 + (1-\alpha)\frac{a - c^{(0)}_1}{K + c_{\text{tot}} - c^{(0)}_1}\right]\right]\,, && (109)\end{aligned}$$

with

$$\omega = \theta \left[ 1 - \frac{K}{\alpha(a - c_1^{(0)}) + K} \right] . \tag{110}$$

We use the fact that

$$c_{\text{atp}} = \alpha c_1^{(0)} + O(\theta) . \tag{111}$$

Thanks to Eq. (111), when considering the value at the midpoint, the approximation

$$\alpha(a - c_1^{(0)*}) \simeq z , \tag{112}$$

holds for every term contained in the expression multiplying $\theta$, because any correction would be of the order of $\theta^2$. Additionally, we have

$$c_{\text{tot}}^* - c_1^{(0)*} = \frac{K c_1^{(0)*}}{a - c_1^{(0)*}} . \tag{113}$$

Using these expressions, we rearrange the formula:

$$c_{\text{atp}}^* = \alpha c_1^{(0)^*} + c_{\text{tot}}^* \omega \left[ 1 - \frac{\frac{K c_1^{(0)*}}{a - c_1^{(0)*}}}{K + (a - c_1^{(0)*}) + \frac{K c_1^{(0)*}}{a - c_1^{(0)*}}} \left[ 1 + (1 - \alpha) \frac{(a - c_1^{(0)*})^2}{Ka} \right] \right] \tag{114}$$

$$= \alpha c_1^{(0)^*} + c_{\text{tot}}^* \omega \left[ 1 - \frac{K c_1^{(0)*}}{K(a - c_1^{(0)*}) + (a - c_1^{(0)*})^2 + K c_1^{(0)*}} \left[ 1 + (1 - \alpha) \frac{(a - c_1^{(0)*})^2}{Ka} \right] \right] \tag{115}$$

$$= \alpha c_1^{(0)^*} + c_{\text{tot}}^* \omega \left[ 1 - \frac{K(\alpha a - z)}{z^2/\alpha + \alpha K a} \left[ 1 + \frac{1 - \alpha}{\alpha^2} \frac{z^2}{Ka} \right] \right] \tag{116}$$

$$= \alpha c_1^{(0)^*} + \omega \left[ a - \frac{z}{\alpha} + \frac{\alpha K a}{z} - K \right] \left[ 1 - \frac{K(\alpha a - z)}{z^2/\alpha + \alpha K a} \left[ 1 + \frac{1 - \alpha}{\alpha^2} \frac{z^2}{Ka} \right] \right] . \tag{117}$$

To solve this equation, we are left with the only term containing $c_1^{(0)*}$ for which we cannot use Eq. (112), and we rather have to use the solution of the mass balance equation for $c$, namely Eq. (31). First, we lighten the notation by defining

$$\begin{aligned}
\xi &:= \omega \left[ a - \frac{z}{\alpha} + \frac{\alpha K a}{z} - K \right] \left[ 1 - \frac{K(\alpha a - z)}{z^2/\alpha + \alpha K a} \left[ 1 + \frac{1 - \alpha}{\alpha^2} \frac{z^2}{Ka} \right] \right] \\
&= \omega \left[ a - \frac{z}{\alpha} + \frac{\alpha K a}{z} - K \right] \left[ 1 - \frac{\alpha K(\alpha a - z)}{z^2 + \alpha^2 K a} \left[ \frac{\alpha^2 K a + (1 - \alpha) z^2}{\alpha^2 K a} \right] \right] \\
&= \omega \left[ a - \frac{z}{\alpha} + \frac{\alpha K a}{z} - K \right] \left[ 1 - \frac{\alpha^2 K a - \alpha K z}{\alpha^2 K a} \frac{\alpha^2 K a + z^2 - \alpha z^2}{\alpha^2 K a + z^2} \right] \\
&= \frac{\omega}{\alpha} \left[ 1 + \frac{\alpha K}{z} \right] [\alpha a - z] \left[ 1 - \frac{\alpha a - z}{\alpha a} \left[ 1 - \alpha \frac{z^2}{z^2 + \alpha^2 K a} \right] \right] .
\end{aligned} \tag{118}$$

We use this definition of $\xi$ to obtain an expression for $c_{\text{tot}}^*$:

$$z = \alpha a - c_{\text{atp}}^* = \alpha a - \alpha \frac{a + c_{\text{tot}}^* + K - \sqrt{(a + c_{\text{tot}}^* + K)^2 - 4 a c_{\text{tot}}^*}}{2} - \xi , \tag{119}$$

and thus

$$(-a + c_{\text{tot}}^* + K)^2 + 4Ka = [2z/\alpha + c_{\text{tot}}^* + K - a + 2\xi/\alpha]^2 \tag{120}$$

$$= 4 \frac{(z + \xi)^2}{\alpha^2} + 4 \frac{z + \xi}{\alpha} (c_{\text{tot}}^* + K - a) + (c_{\text{tot}}^* + K - a)^2 . \tag{121}$$

By solving for $c_{\text{tot}}^*$ we have

$$\begin{aligned}
c_{\text{tot}}^* &= \frac{\alpha^2 K a - (z + \xi)^2}{\alpha(z + \xi)} + a - K \\
&= \frac{(\alpha a - (z + \xi))(\alpha K + (z + \xi))}{\alpha(z + \xi)} .
\end{aligned} \tag{122}$$

Note that when $\xi = 0$ we recover the formula obtained previously for $c^*_{\text{tot}}$.

The volume at initiation is

$$V^* = \frac{n^*}{c^*_{\text{tot}}} = \frac{n^*\alpha(z-\xi)}{(\alpha a - (z+\xi))(\alpha K + (z+\xi))} \simeq \frac{n^*\alpha}{\alpha a - (z+\xi)} \,. \tag{123}$$

We expand for small $\xi$ to obtain

$$V^* \simeq \frac{n^*\alpha}{\alpha a - z} + \frac{n^*\alpha}{(\alpha a - z)^2}\xi \,. \tag{124}$$

The contribution from the special binding sites is

$$f(\alpha) := \frac{n^*\alpha}{(\alpha a - z)^2}\xi = \frac{n^*}{(\alpha a - z)}\omega \left[1 + \frac{\alpha K}{z}\right]\left[1 - \frac{\alpha a - z}{\alpha a}\left[1 - \alpha\frac{z^2}{z^2 + \alpha^2 K a}\right]\right] \,. \tag{125}$$

The function in Eq. (125) is a monotonically decreasing function of $\alpha$, showing that the effect of deactivation is enhanced by the special binding sites.

### 5 Derivation of the fraction of active DnaA from a mechanistic model

The process by which DnaA is activated and de-activated through reactions of ATP hydrolysis and nucleotide exchange has been studied extensively. The known regulatory mechanisms are RIDA and datA for the hydrolysis of ATP, and DARS1/DARS2 and the membrane phospholipids for the exchange from ADP to ATP. However, it was shown [2, 3, 4] that the system can function even without datA and DARS1/DARS2, that play a supporting role. In this section we derive the fraction of acive DnaA under the simplifying assumption that RIDA is the only active regulatory mechanism involved in the interconversion between ATP and ADP. Additionally, a DnaA-ATP molecule can spontaneously convert to DnaA-ADP at a lower rate. The main factor restoring the active state is assumed to be the synthesis of new DnaA proteins, that are more likely to bind ATP than ADP, because ATP is more abundant.

The main characteristic of RIDA is that the hydrolysis is related with the presence of the replication fork. Consequently, previous models [1, 5] incorporate hydrolysis via Michaelis Menten equation that depends the number of replication forks $n_{\text{forks}}$:

$$\text{hydrolisis rate} = k_{\text{forks}}\frac{n_{\text{forks}}}{V}\frac{a_{\text{atp}}}{a_{\text{atp}} + K_{\text{forks}}} \approx k_{\text{forks}}\frac{n_{\text{forks}}}{V} \,. \tag{126}$$

where $k_{\text{forks}}$ and $K_{\text{forks}}$ are constants, and the approximate equality descends from the assumption that RIDA operates near saturation. The corresponding dynamical equation for the concentration of active DnaA is

$$\frac{da_{\text{atp}}}{dt} = -k_{\text{forks}}\frac{n_{\text{forks}}}{V} - a_{\text{atp}}k_h + (\Lambda + k_e)a_{\text{adp}} \,, \tag{127}$$

where $k_h$ and $k_e$ are the intrinsic rates of hydrolysis and synthesis, respectively, and $\Lambda$ is the growth rate. The equation is coupled with the conservation law $a_{\text{atp}} + a_{\text{adp}} = a$. At steady state, we have

$$k_{\text{forks}}\frac{n_{\text{forks}}}{V} + a_{\text{atp}}k_h = (\Lambda + k_e)(a - a_{\text{atp}}) \,, \tag{128}$$

and thus

$$\alpha = \frac{\Lambda + k_e}{k_h + k_e + \Lambda}\left[1 - \frac{k_{\text{forks}}n_{\text{forks}}}{Va(\Lambda + k_e)}\right] \,. \tag{129}$$

Defining

$$\alpha_{\max} = \frac{\Lambda + k_e}{k_h + \Lambda + k_e} \,,$$
$$\chi_0 = \frac{k_{\text{forks}}}{a(\Lambda + k_e)} \,, \tag{130}$$

we obtain the phenomenological relation in the Main Text for $\alpha > \alpha_{\min}$:

$$\alpha = \alpha_{\max}\left[1 - \frac{\chi_0 n_{\text{forks}}}{V}\right] \,. \tag{131}$$

Considering a zero-th order reaction for the hydrolysis ($a_{\text{atp}} \gg K_{\text{forks}}$) is no longer possible as $\alpha$ approaches zero. A comparison with the curve without approximation (Fig.3) shows that the deviation is only relevant far from the volume at initiation in fast growth conditions in case $K_{\text{forks}} = 10\,nM (> 1\mu m^3$, see [6]).

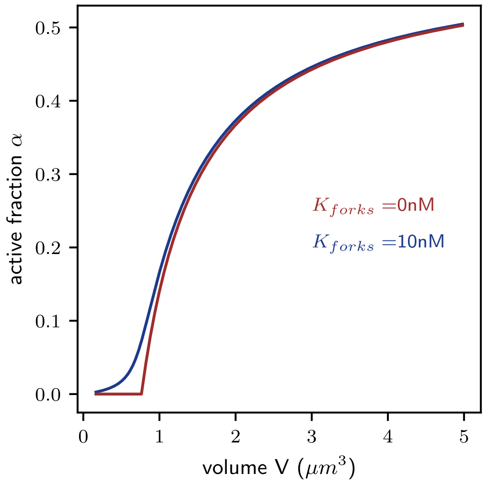

Figure 3: Comparison of the steady state solution of the equation for $\alpha$ in the case a Michaelis Menten kinetics (blue) or a zeroth order reaction kinetics (red) are used.
